# TRPML3 regulates neuronal gene expression in an *in vitro* model of autophagy and may act as a genetic marker of familial neurodegenerative disorders

**DOI:** 10.1101/2025.06.28.662125

**Authors:** Sushanth Adusumilli, Mahesh Mahadeo Mathe, Jayasha Shandilya, Tapan Kumar Nayak

## Abstract

Autophagy is a conserved pro-survival pathway for delivering misfolded proteins and damaged organelles to lysosomes for degradation and protein homeostasis. Anomaly in autophagy leads to aberrant protein aggregation in neuronal cells, which is a common etiology of neurodegenerative disorders. Endo-lysosomal cation channel TRPML3 (Transient Receptor Potential Mucolipin-3) has been shown to induce autophagy in cell line models. However, the mechanism of TRPML3 mediated autophagy induction and the underlying gene expression changes are not clearly understood. Here, by using Ca^2+^-and electrical-current measurements, RNA sequencing and RT PCR studies, we explored the cellular function of TRPML3 and the global transcriptomic profile in a cell-based serum starvation model of autophagy. We report that serum starvation leads to downregulation of neuronal developmental genes during autophagy induction. TRPML3 overexpression further amplifies the effect of starvation in downregulating neuronal gene expression. But, when nutrition is not a limiting condition, TRPML3 overexpression upregulated neuronal genes including those responsible for axon guidance, synaptogenesis, and dendritic arborization. TRPML3 mediated neuronal gene expression changes were, presumably, due to transcription factors (TF) TFEB, FOXO1 and neuron-specific TFs such as SOX2, and ETV5. To further validate the role of TRPML3 in neuronal gene regulation, we performed meta-analysis of publicly available RNAseq datasets on neurodegenerative disorders which provided insight into the heterogeneity in the molecular mechanisms of autophagy and corroborated the TFEB-mediated autophagy induction and neuronal gene expression in TRPML3 overexpression condition. Based on our results, we propose that TRPML3 may act as a potential genetic marker for familial neurodegenerative disorders.

## INTRODUCTION

Neurodegenerative disorders are globally one of the leading causes of deaths and disability-adjusted life years (DALY: years lived with disability), thus imposing significant public health and socio-economic burden (1, 2). Most of the neurodegenerative disorders are sporadic or idiopathic in nature though a significant percentage of them are genetic with Mendelian inheritance patterns (1). A common etiology of most of the neurodegenerative disorders, both sporadic and familial, such as Parkinson’s, Alzheimer’s, Tauopathies, ALS, Huntington’s, Cockayne syndrome (3) and Ataxia-Telangiectasia (4), is aggregation of misfolded proteins. Autophagy (self-devouring) is an evolutionarily conserved cellular process that promotes degradation of misfolded proteins and damaged organelles (5, 6). Therefore, anomaly in autophagy pathways has been strongly implicated in the severity of neurodegenerative disorders (7, 8). Suppression of autophagy has been shown to accelerate aging and neurodegeneration (9). On the flip side, enhancing autophagy flux is considered as a potential therapeutic approach for delaying or preventing neurodegeneration in aged population (10).

Formation of autophagosomes, their shuttling through the common endo-lysosomal pathway, and fusion with the lysosomes for degradation of their contents (a.k.a. autophagy flux) is tightly regulated by multiple protein complexes encoded by autophagy related genes (ATGs) (10–12). Mice deficient in ATG5 (13) and ATG7 (14) in the neuronal cells show progressive accumulation of misfolded proteins, formation of inclusion bodies and eventually neurodegeneration. Mutations in some of the ATGs, including PTEN-induced kinase1 (PINK1) and PARKIN, result in Parkinson’s disease whereas mutations in PRESENILIN 1 lead to Alzheimer’s disease (8). Further, functional deficiency of AMBRA1 (Activating Molecule in Beclin1-Regulated Autophagy) in mouse embryo is associated with accumulation of ubiquitinated proteins and neural tube defects (15). So far, more than 600 ATGs have been identified which function as: 1. core autophagy, 2. autophagy regulators, 3. MTOR and upstream pathways, 4. mitophagy (autophagy of mitochondria), 5. docking and fusion, and, 6. lysosome and lysosome related genes (12). However, there may be other potential ATGs and fringe regulators of autophagy, which are yet to be annotated. Further, there is a great deal of biological heterogeneity in ascribing function to these ATGs depending on the model systems (human or animal tissue, induced pluripotent stem cell (iPSC) derived or immortalized cell lines), variations in experimental methods and technical handling. Therefore, a comprehensive investigation of the role of these genes in the autophagy pathway in the context of neurodegenerative disorders is highly warranted.

Ion channels are critical cell membrane proteins which conduct ions, and generate electrical currents, and (16) are responsible for sensory perception, brain computing, muscle contraction, cardiac rhythm, and immune response. They are implicated in several human disorders, including Alzheimer’s, Parkinson’s, and Schizophrenia (17). Ion channels expressed in the intracellular organelle membranes (18) maintain ionic homeostasis, [Ca^2+^] and pH balance and are involved in cellular cargo transport, metabolism, and cell signaling. Some of these ion channels, such as ER-resident Ca^2+^ channels, IPTR and RYR, lysosomal two-pore channel gene (TPCN) and mucolipin family TRP channel genes (TRPML) have been implicated in the regulation of autophagy flux (19) though there are conflicting reports about their function in autophagy (20). The TRPML family, comprising of TRPML1, TRPML2 and TRPML3 genes, express in the endo-lysosomal compartments, regulate endocytosis and autophagy flux (21–23). Loss-of-function of TRPML1 causes lipid storage disorder, type-IV mucolipidosis (MLIV), a neurodegenerative disease that causes mental retardation and retinal degeneration (24, 25). Gain-of-function in TRPML3 gene is implicated in *varitint-waddler* (*Va*) phenotype with hearing and pigmentation defects (26–28).

TRPML3 expression in the membrane and intracellular compartments is known to be dynamic, with recruitment of TRPML3 to autophagosomes upon induction of autophagy (29). Accordingly, overexpression of TRPML3 leads to increased autophagy and is known to play a role in evoked autophagy by cell stressors such as starvation (30, 31). Further, TRPML3 has been shown to provide Ca^+2^ for membrane fusion during autophagosome formation (31) and autophagosome-lysosome fusion (32). TRPML3 is the only ion channel that is expressed in the autophagosomes (19) and, like TRPML1, has significant expression in the late endosomes/lysosomes (29, 33).

However, TRPML3 is known to get inactivated by low pH in the lysosomes (34). Therefore, how TRPML3 functions in concert with other ion channels and ATGs to orchestrate the autophagy flux is still not clearly understood.

Here, in an *in vitro* serum starvation model of autophagy, we show the significance of a TRPML3 mediated Ca^2+^ response in the regulation of autophagy flux by using a combination of fluorescence imaging, patch-clamp electrophysiology, and calcium recordings. Our transcriptomic studies indicate that TRPML3 overexpression leads to differential regulation of some of the key transcription factors, such as TFEB, FOXO3 and FOXO1 and enrichment of key neuronal gene sets (in the HEK293 cells) which are implicated in synaptogenesis, axon guidance, spine morphogenesis and neurotransmitter mechanism. Further, by using an elaborate meta-analysis of RNA sequence data from more than 30 neurodegenerative disorders, we identified a small subset of key endo-lysosomal ion channels including TRPML3, ATGs (ATG7 and ATG9B) and transcription factors (TFEB, FOXO3, FOXO1, ETVs) which are critical for autophagy flux and are differentially regulated in idiopathic vs familial neurodegenerative disorders. Overall, our results suggest that TRPML3 plays a key role in autophagy and can potentially be a genetic denominator in familial neurodegenerative disorders.

## RESULTS

### TRPML3 overexpression and activation induces autophagy

The ion channel function of TRPML3 has been recently implicated in autophagy induction (31, 32, 35). To quantitatively estimate the effect of TRPML3 expression/activation on autophagy flux, we heterologously expressed it in an *in vitro* serum starvation model of autophagy (see methods). In total, we had 4 experimental conditions: 1. Control vector transfected cells with 10% serum (control Fed or V), 2. Control vector transfected cells without serum (control starved or VS), 3. Cells over-expressing TRPML3 in 10% serum (ML3 Fed or T), and 4. TRPML3 transfected cells without serum (ML3 starved or TS). Fig. 1a and 1b show cells expressing LC3-EGFP (microtubule-associated protein 1 light chain 3; a marker of autophagy induction (36)) under control Fed (Fig. 1a) vs control starved conditions (Fig. 1b), respectively. We counted the number of puncta to quantify autophagy induction in the cells expressing specific markers of autophagy. Fig. 1g are bar plots showing the number of LC3 (left) and TRPML3 (right) puncta under different experimental conditions expressed as mean ± S.E.M. Serum-starvation significantly increased the number of LC3 puncta (17 ± 3, Fig. 1b) vs the fed condition (6 ± 1, Fig. 1a), which showed autophagy induction by serum starvation. Fig. 1c and 1d show the representative confocal images of HEK293 cells expressing LC3-EGFP/TRPML3-mCherry under fed (TRPML3 Fed or T) or serum-starvation conditions (TRPML3 starved or TS). When TRPML3 was co-expressed, the number of LC3 puncta was significantly higher in fed condition (12±3, Fig. 1c) vs LC3 overexpressed fed condition (6±1). It indicated that TRPML3 over-expression exaggerated the flux of autophagy. Starvation does not significantly increase the number of LC3 puncta (15±3, Fig. 1d) under TRPML3 overexpression. On the other hand, starvation significantly increased the number of TRPML3 puncta (52±9, Fig. 1d) vs fed condition (22±5, Fig. 1c), which suggested intracellular mobilization of TRPML3 under evoked autophagy.

**Fig. 1.**
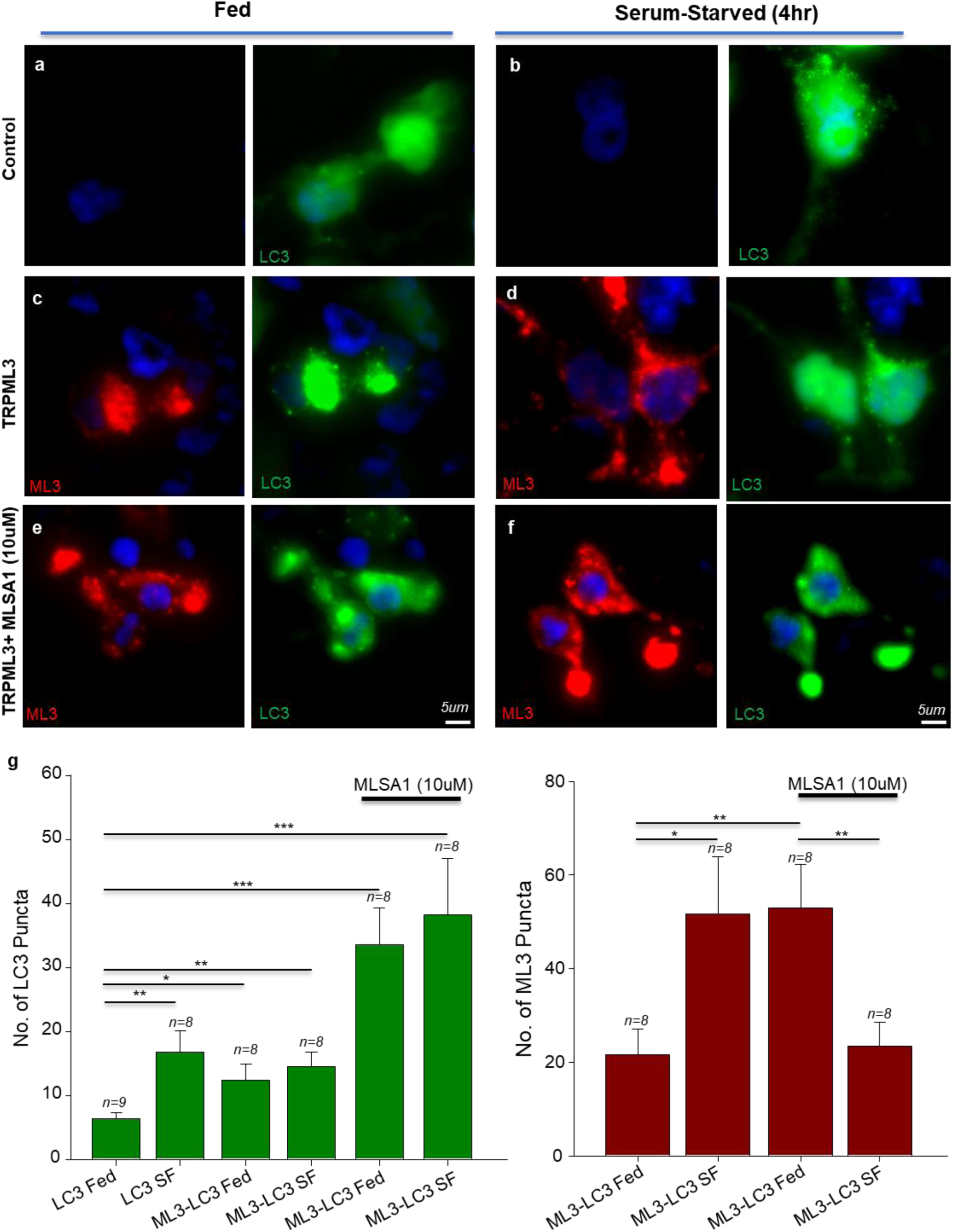
TRPML3 overexpression induces autophagy: Representative zoomed maximum intensity projections of Z-stack images of HEK293T cells under the following experimental conditions: **(a, b)** LC3-EGFP overexpression under fed (a) or serum-starved conditions (b); **(c, d)** TRPML3-mCherry and LC3-EGFP overexpression under fed (c) or serum-starved conditions (d); **(e, f)** TRPML3-mCherry and LC3-EGFP overexpressed conditions and treatment with 10μM MLSA1 under fed (e) or serum-starved conditions (f). Note the punctate autophagolysosome structures under serum starvation and TRPML3 over-expression conditions **(g)** Bar plots representing the average number of LC3 (left) and TRPML3 (right) puncta observed under the mentioned conditions. *n= number of Field of Views (FoVs) for each condition. * = 0.01<p<0.05, ** = 0.005<p<0.01, *** = p<0.005*

Further, we confirmed the ion channel function of TRPML3 in autophagy induction by treating the cells with 10μM of MLSA1, a TRPML non-specific agonist. In cells co-expressing LC3 and TRPML3, the number of LC3 puncta under fed condition increased by 5-fold (34 ± 5, Fig. 1e) compared to LC3 overexpressed fed condition when treated with 10μM MLSA1, which further corroborated that TRPML3 over-expression exaggerated autophagy flux. Starvation along with MLSA1 treatment further increased the number of LC3 puncta (39 ± 7, Fig. 1f). MLSA1 treatment led to a significant increase in the number of TRPML3 puncta in fed condition (53 ± 9, Fig. 1e). However, MLSA1 treatment during starvation significantly decreased the number of TRPML3 puncta (23±6, Fig. 1f) vs the fed condition (Fig. 1e), presumably due to fusion of TRPML3 containing vesicles with lysosomes. Though MLSA1 treatment significantly decreased the number of TRPML3 puncta, in those cells, the number of LC3 labeled puncta were higher indicating to significant increase in the number of puncta corresponding to autophagosome formation.

Formation and elongation of autophagosomes typically happen on the ER-mitochondria (37, 38), ER-Golgi intermediate compartment (39) or the plasma membrane (40, 41). Autophagosomes containing the desired cargo then shuttle through the common endo-lysosomal pathway for degradation of their contents. Several ATGs play specific roles throughout the autophagy pathway. To understand the role of TRPML3 in autophagy pathway, we studied its co-localization with autophagy markers during autophagy induction and progression. Fig. 2a-h are representative confocal images of HEK293 cells fluorescently co-labelled with TRPML3 and specific markers for autophagy-induction (ATG7, and LC3A/B) and markers for the sites of autophagy initiation in the intracellular organelles (KDEL, lysotracker) under fed or serum-starvation conditions.

**Fig. 2.**
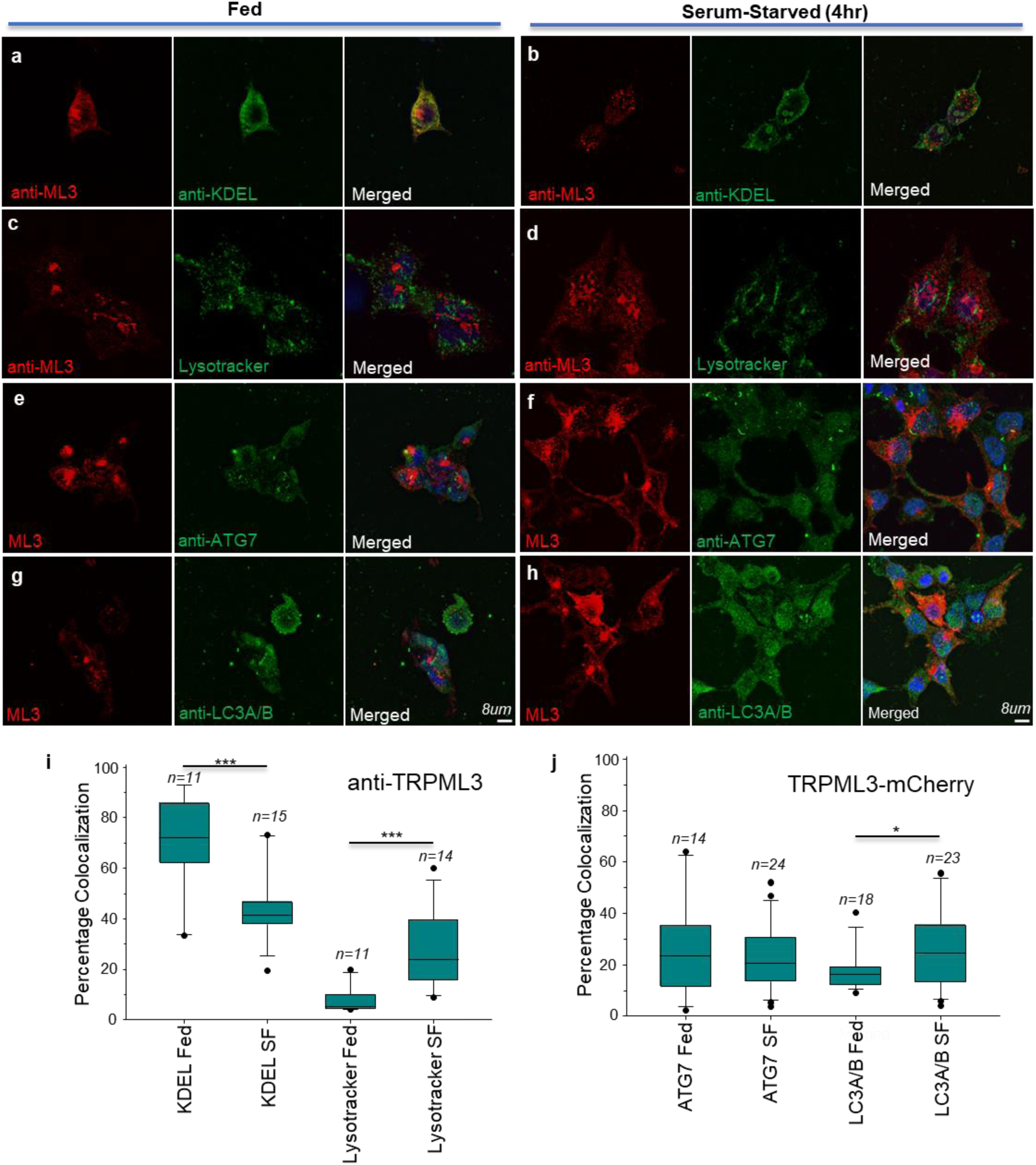
TRPML3 colocalizes with key autophagy markers: Representative maximum intensity projections of confocal Z-stack images of HEK293T cells under the following conditions: **(a, b)** Cells stained with anti-TRPML3 and anti-KDEL antibodies under fed (a) vs serum-starved (b) conditions. **(c, d)** Cells stained with anti-TRPML3 antibody and loaded with 100nM Lysotracker Green under fed (c) vs serum-starved conditions (d). **(e, f)** Cells overexpressing TRPML3-mCherry and stained with anti-ATG7 antibody under fed (e) or serum-starved conditions (f). (**g, h)** Cells overexpressing TRPML3-mCherry and stained with anti-LC3A/B antibody under fed (g) or serum-starved conditions (h). **(i, j)** Box plots showing the Manders’ colocalization coefficients, converted to percentage, of anti-TRPML3 antibody (i) and TRPML3-mCherry (j) with denoted markers. Black dots represent the outliers for each box. *n = number of Region of Interests (RoIs) for each condition. * = p<0.05, *** = p<0.005*

Boxplots in Fig. 2i and 2j show the Manders colocalization coefficients determined from the analysis of the images shown in Fig.2a-2h. It was observed that the endogenous TRPML3 shows significantly higher % colocalization with KDEL (Endoplasmic Reticulum marker) in the fed (Median, IQR; 72%, 62-85%, Fig. 2a) vs the starved conditions (Median, IQR; 40%, 37-45%, Fig. 2b). On the other hand, the endogenous TRPML3 shows negligible colocalization with lysotracker (Late Endo-lysosomal marker) in the fed (Median, IQR; 5%, 4-9%, Fig. 2c) vs starved conditions (Median, IQR; 24%, 15-39%, Fig. 2d). Further, overexpressed TRPML3 showed significantly higher colocalization with LC3A/B (marker for autophagosomes) in starved conditions (Median, IQR; 25%, 14-36%, Fig. 2h) vs fed conditions (Median, IQR, 15%, 11-18%, Fig. 2g), while it showed no significant changes in colocalization with ATG7 (Fig. 2e,2f), a core ATG involved in autophagosome biogenesis.

### Ca^+^ intrusion through TRPML3 induces autophagy

To study the functional role of TRPML3 channels during autophagy induction, we performed calcium signaling studies and whole-cell patch clamp electrophysiology experiments. Fig. 3a shows the representative ΔF/F_0_ values for the cytosolic [Ca^2+^]_i_ (change in [Ca^2+^]) in control vs TRPML3 overexpressed HEK293 cells in the presence of 5μM MLSA1 under fed or serum-starved conditions. Fig. 3c and 3d are bar plots that show the peak [Ca^2+^]_i_ (ΔF/F_0_) and total [Ca^2+^]_i_ (ΔA/A_0_) released, respectively, in HEK cells under the above conditions in the presence of MLSA1 treatment. Serum starvation led to significant increase in the peak and total [Ca^2+^]_i_ in both the control and TRPML3 overexpressed HEK cells (Mean ± S.E.M; Ca^2+^ peak: control Fed: 0.04±0.005 vs control starved: 0.32±0.03, TRPML3 Fed: 0.1±0.005 vs TRPML3 starved: 0.85±0.04 and Ca^2+^ total: Ca^2+^ peak: control Fed: 0.05±0.005 vs control starved: 0.24±0.04, TRPML3 Fed: 0.1±0.005 vs TRPML3 starved: 0.73±0.05). We observed that serum starvation induced ∼ 5-8-fold increase in [Ca^2+^]_i_ response in MLSA1 treated cells under fed vs starved conditions, whereas TRPML3 overexpression caused ∼ 3-fold increase in [Ca^2+^]_i_ response under fed vs starved conditions. Table 1 shows the Mean ± S.E.M of time constant (τ_rise_) of exponential rise to the maximum for [Ca^2+^]_i_ response under the different experimental conditions. Serum starvation led to a slow, but, sustained [Ca^2+^]_i_ loading (see Fig. 3a) vs the fed conditions (τ_rise_ in sec: control Fed: 5.9 ± 0.2 and TRPML3 Fed: 4.6 ± 0.1 vs control starved: 30.2 ± 0.3 and TRPML3 starved: 9.2 ± 0.1). But TRPML3 over-expression, in fed vs serum starved conditions respectively, showed 1.5 to 3-fold faster [Ca^2+^]_i_ rise to maximum vs the corresponding untransfected cells. Further, to differentiate the contribution of external vs internal Ca^2+^ to the cell [Ca^2+^]_i_, we performed Ca^2+^ recordings in the presence of EGTA, a Ca^2+^ chelator (see Fig. 3a and Fig. S1). The peak [Ca^2+^]_i_ rise and the total cell [Ca^2+^]_i_ in the presence of EGTA was small (∼50%) compared to control fed condition, suggesting that in our *in vitro* serum starvation model most of the Ca^2+^ response was primarily due to the TRPML3 expressed on the cell membrane. Fig. S1 shows the Ca^2+^ response in the presence of EGTA under control and TRPML3 over-expressed conditions which were small but detectable and followed similar kinetics (Table 1).

**Fig. 3.**
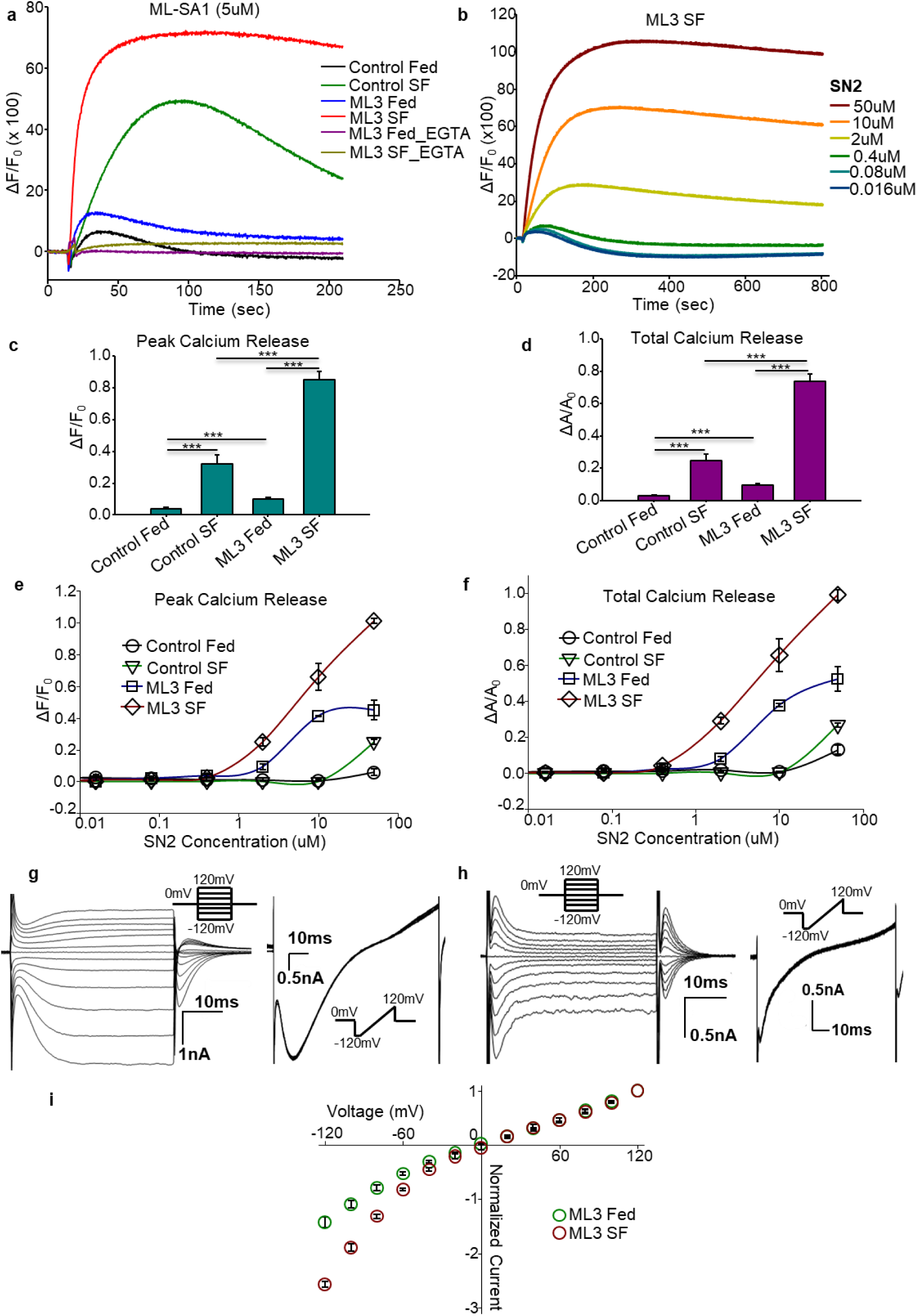
Calcium currents through TRPML3 channels induce autophagy: **(a)** Representative [Ca]_i_ recordings showing ΔF/F0 changes vs time from HEK293T cells under different conditions in the presence of 5μM MLSA1. **(b)** Representative [Ca]_i_ recordings from serum-starved HEK293T cells overexpressing TRPML3 in the presence of different [SN2], a TRPML3 specific agonist. **(c)** Bar plot showing peak [Ca]_i_ from the given experimental conditions upon MLSA1 treatment. **(d)** Bar plot showing the total [Ca]_i_ from the shown experimental conditions upon MLSA1 treatment. For MLSA1 treatment experiments without EGTA, data was acquired from at least 20 wells of cells for each experimental condition. For experiments with EGTA, data was acquired from 6 wells of cells. **(e)** Line and Scatter plot showing peak [Ca]_i_ from the denoted experimental conditions in the presence of different [SN2]. **(f)** Line and Scatter plot showing total [Ca]_i_ from the denoted experimental conditions in the presence of different [SN2]. For SN2 treatment experiments, data was acquired from 3 wells of cells in each experimental condition. **(g, h)** Representative whole cell current recordings from fed (g) and serum-starved HEK293T cells overexpressing TRPML3 on the application of step (left) and ramp membrane voltage pulse protocols (right), respectively. The pulse protocols are shown in the figures. **(i)** Scatterplot showing the normalized current-voltage (I-V) relationship of TRPML3 under fed or serum-starved conditions. *** = *p < 0.005*.

**Table 1.** Average ± S.E.M of time constant of Calcium release kinetics under the labelled experimental conditions.

| Time Constant<br>(sec) | **Control**<br>**Fed** | **Control**<br>**SF** | **ML3**<br>**Fed** | **ML3**<br>**SF** |
| --- | --- | --- | --- | --- |
| No EGTA | 5.9 ± 0.2 | 30.2 ± 0.3 | 4.6 ± 0.1 | 9.2 ± 0.1 |
| EGTA |  | 36.5 ± 0.4 |  | 16.3 ± 0.3 |

Additionally, we performed Ca^2+^ recordings with SN2, a TRPML3 specific agonist, under different concentrations. Fig. 3b shows the representative ΔF/F_0_ values of cytosolic [Ca^2+^]_i_ from TRPML3 overexpressed HEK cells under starvation conditions upon treatment with increasing concentrations of SN2. Figs. 3e and 3f are line and scatter plots that show the peak [Ca^2+^]_i_ (ΔF/F_0_) and total [Ca^2+^]_i_ (ΔA/A_0_) respectively at different concentrations of SN2 under different experimental conditions. While there was negligible peak [Ca^2+^]_i_ (ΔF/F_0_) and total [Ca^2+^]_i_ in control HEK cells at [SN2] <50 μM, TRPML3 overexpressed cells showed SN2 mediated calcium response at [SN2] as low as 0.4μM and showed a strong concentration dependent increase in [Ca^2+^]_i_. In the presence of SN2, clearly, serum starvation resulted in at least 2-fold higher Ca^2+^ response vs the fed condition, as observed above with MLSA1, in both the control (Mean ± S.E.M; peak: control Fed: 0.05 vs control starved: 0.2 at 50μM) as well as TRPML3 overexpressed cells (peak: TRPML3 Fed: 0.4 vs TRPML3 starved: 1.0 at 50μM).

To further verify that the observed [Ca^2+^]_i_ response was due to calcium intrusion through the TRPML3 channels, we recorded whole-cell currents from HEK293T cells over-expressing TRPML3 channels. Fig. 3g and 3h show representative whole-cell current traces obtained by applying step (left) or ramp voltage pulse protocol (right) under fed vs serum-starved conditions. The macroscopic currents from serum starved cells (Fig. 3h) were substantially larger and showed greater inward rectification vs the fed condition (Fig. 3g). Fig. 3i shows a scatterplot of normalized current-voltage (I-V) relationship of TRPML3 under fed vs starved conditions which showed significantly larger inward-rectifying currents under serum starvation conditions. Larger inwardly rectifying TRPML3 currents are accompanied by proportionately larger Ca^2+^ flux into the cells.

### TRPML3 regulates the expression of neuronal gene sets in an *in vitro* model of autophagy

TRPML3 current densities and cellular localization were altered in serum-starved cells during autophagy induction, presumably due to change in gene expression of the channel. We posited that the altered expression of TRPML3 may result in a global change in the transcriptomic profile during serum starvation. To understand the potential change in gene expression of TRPML3 and related genes, we performed RNA sequencing (RNAseq) and quantitative RT-PCR studies in the *in vitro* cell-line model in the fed vs serum-starved conditions.

Extended serum starvation leads to significant cell-cycle arrest (42–44). To investigate if the 4 hrs of serum starvation regime in our model led to any significant cell cycle arrest, we, first, performed flow cytometry analyses (see Methods) in HEK cells during serum starvation. Fig 4a shows the distribution of cell populations in different cell cycle phases (G1-S-G2-M) under different conditions. Fig 4b is a bar plot showing the Mean ± S.E.M percentage of cells in each phase. It shows that 4hrs of serum starvation does not cause significant cell cycle arrest. Nocodazole (Noc), an antineoplastic agent, is known to cause cell-cycle arrest (45) at G2/M interface. Fig. 4a (right) shows the effect of progressively increasing [Noc] (75 and 100 ng/ml) on the cell populations. With increasing [Noc], G1, and S phase cell populations decreased and at 100 ng/ml [Noc], most of the cells were observed to be arrested in the G2-M interface. The change in population distribution in the presence of Noc is shown in Fig. 4b. This indicated that 4hr serum starvation had negligible effect on the population distribution insofar as their cell-cycle phases. Noc treatment simply shifted the naturally distributed populations to the G2/M interface.

**Fig. 4.**
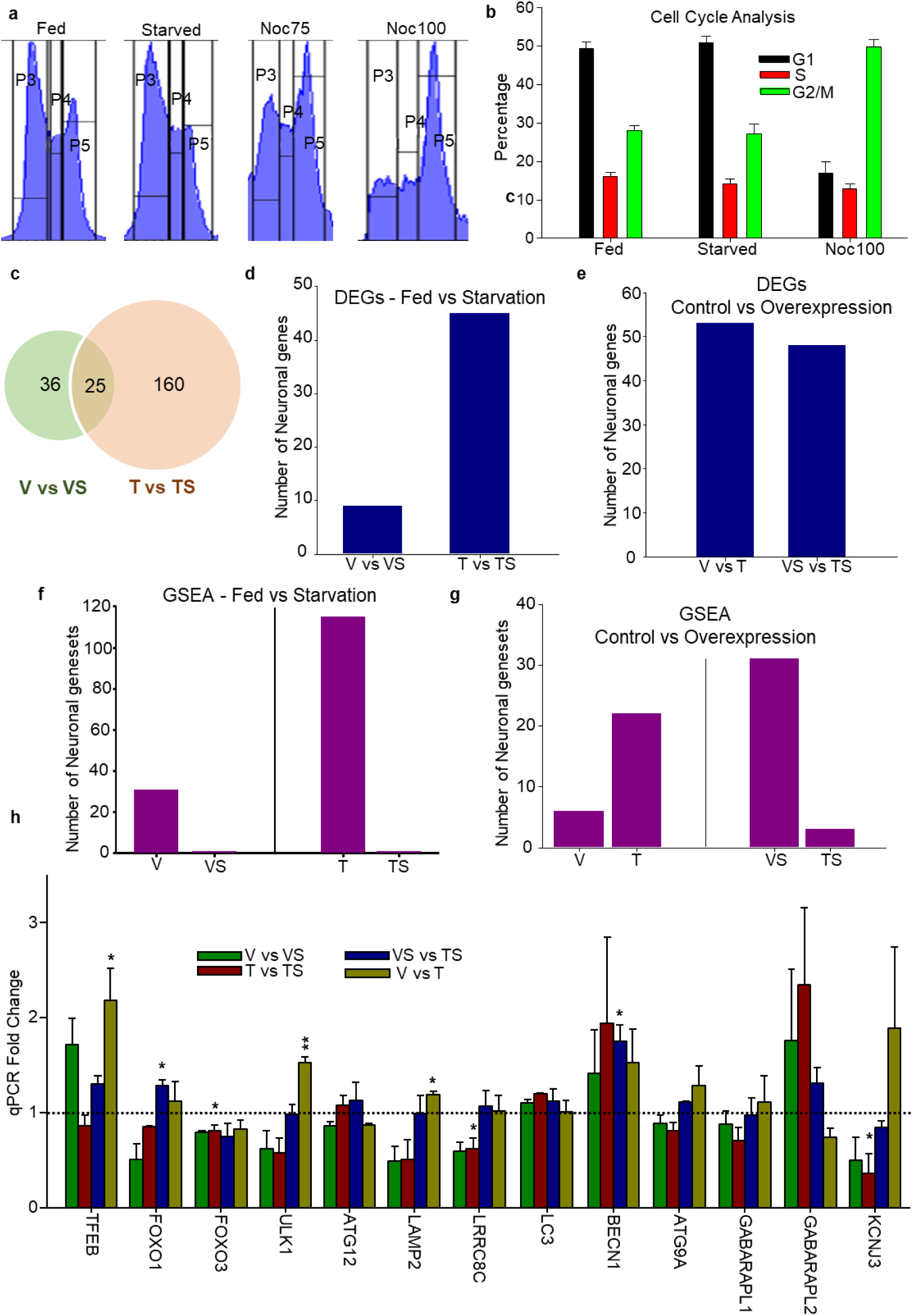
TRPML3 regulates neuronal gene expression in the Starvation Model of Autophagy: (a) Representative flow cytometry data on cell populations in various cell cycle phases under labelled conditions. Noc75/100 – Nocodazole 75ng/ml and 100ng/ml, respectively. P3, P4, and P5 are populations of cells present in G1, S and G2/M phases respectively. Note that the cell population distributions under Fed and starved conditions were comparable, whereas they significantly changed in the presence of Nocodazole. **(b)** Bar plot showing % population of cells in G1/S/ G2/M phases of cell cycle from data in (a). **(c)** Venn diagram representing the total number of Differentially Expressed Genes (DEGs) in the Fed vs Starvation conditions obtained from RNA sequencing analysis. **(d,e)** Bar plots showing the number of neuronal DEGs in Fed vs Starvation (d) and Control vs Overexpression (e) comparisons. **(f, g)** Bar plots showing the number of enriched neuronal gene sets from Gene Set Enrichment Analysis (GSEA) in 4 different pairwise comparisons. Fed vs Starvation (f) represented by ‘V vs VS’ and ‘T vs TS’ comparisons. Ctrl vs Overexpression (g) represented by ‘V vs T’ and ‘VS vs TS’ comparisons. **(h)** Bar plot showing the qPCR fold changes of multiple ATGs and ion channels in denoted pairwise comparisons. Data is acquired from at least three biological replicates. ** = 0.005 < p < 0.01, * = 0.01 < p < 0.05.

### RNAseq studies to understand the role of TRPML3 in autophagy induced by starvation

For RNA-seq and RT-qPCR experiments to quantify gene expressions, we isolated mRNA from cell populations under the following conditions: vector fed (V), vector starved (VS), TRPML3-Fed (T), and TRPML3-Starved (TS). Further, we performed differential Gene Expression (DGE) studies using two different pipelines (*Stringtie-DESeq2* and *FeatureCounts-DESeq2*) and Gene Set Enrichment Analysis (GSEA) as described in the methods (Fig S2a).

Fig. 4c is a Venn diagram that shows the number of total DEGs (average from both pipelines) in *T vs TS* and *V vs VS* comparisons. Both the pipelines predicted substantially higher numbers of unique DEGs in the *T vs TS* comparison versus the *V vs VS* comparison (Fig 4c). As many as 3 times higher numbers of genes were predicted to be differentially expressed upon starvation in TRPML3 overexpressed conditions as opposed to the control. Further, most of the genes that were differentially expressed in TRPML3 overexpression background were neuron specific or had neuronal functions. Fig. 4d and 4e show the number of unique neuronal genes (average from both pipelines) that were differentially expressed due to starvation (Fig 4d, *V vs VS* and *T vs TS* comparisons) and TRPML3 overexpression (Fig 4e, *V vs T* and *VS vs TS* comparisons). *T vs TS* comparison shows significantly higher number of neuronal DEGs (45 genes) compared to *V vs VS* comparison (10 genes, Fig 4c). However, there is no significant change in the number of neuronal DEGs between *V vs T* and *VS vs TS* comparisons (Fig 4d). Table S1 shows selected neuronal DEGs and their expression patterns that were called significant by at least one of the RNA-seq pipelines in different pairwise comparisons. The DEGs were broadly grouped into 3 categories as shown in Table S1: Transcription factors (TFs), structural and signaling proteins, and genes with reported mutations that cause neurological disorders (e.g. C9orf72 [in ALS and FTD], CLIC2 [in intellectual development disorder], and SETX [spinocerebellar ataxia with axonal neuropathy]; OMIM database). Of these, ETV5, and ETV1 are key TFs expressed in multipotent neural crest cells (46) and developing synapses (47) respectively, where they play key roles in regulating transcriptional activation and synaptic maturation of NMDA and GABA receptors. SOX2 is a global transcription factor that regulates gene expression in neural progenitor cells and are implicated in a spectrum of CNS defects, including vision and hippocampus impairments, intellectual disability, and motor control problems (48, 49). Further, GDF7 and FOXC1 are critical transcription factors which are linked to development of the spinal cord (50) and cerebellum (51). Majority of the TFs and genes with disease-causing mutations were downregulated in *T vs TS* comparison while most of the structural and signaling proteins were upregulated in *T vs TS* and *V vs T* comparisons (Table S1).

Gene set enrichment analysis (GSEA) showed the effects of serum starvation (Fig. 4f) and TRPML3 overexpression (Fig. 4g) on various molecular pathways and biological functions in the model system. The neuronal gene sets were downregulated in serum-free conditions (Fig. 4f, left). The effect of serum starvation was more pronounced in the TRPML3 over-expression background with more than 3-fold enrichment in *T vs TS* over *V vs VS* conditions (Fig. 4f, right). This was further corroborated by the GSEA analysis of the *VS vs TS* comparisons (Fig. 4g, right). Tables S2 and S3 show the list of enriched neuronal gene sets common to both pipelines in specific pairwise comparisons. Most of them are associated with neuronal development and function, sensory perception, synaptic organization, and neurotransmitter release. Multiple autophagy-related gene sets were found to be enriched in starvation conditions as expected (Table S4).

Overexpression of TRPML3 was further validated using RT-qPCR (Fig S2b). Fig 4h is a bar plot showing qPCR fold changes of crucial ATGs/ARGs and ion channels involved in the canonical autophagy pathway. Overexpression of TRPML3 under fed conditions (*V vs T*) leads to significant upregulation of TFEB, a master regulator of lysosomal biogenesis (52–54). Further, ULK1 and LAMP2 are also significantly upregulated under the same conditions. Transcription factors FOXO1 and FOXO3 are respectively upregulated (*VS vs TS*) and downregulated (*T vs TS*) due to TRPML3 overexpression under starvation conditions. Both are reported to have crucial roles in neuronal development and autophagy (55–57). Other important genes include BECN1 (upregulation in *VS vs TS*), ion channels LRRC8C and KCNJ3 (downregulation in *T vs TS*). Overall, TRPML3 overexpression directly enhanced autophagy and lysosomal function by regulating TFEB, FOXO1 and other autophagy related genes.

### TRPML3 as a potential marker for familial neurodegenerative disorders

TRPML3 profoundly altered the gene expression profile of specific genes involved in neuronal development, synapse formation etc. in the cultured cells under serum starvation conditions during autophagy induction. This posed a titillating question if TRPML3 plays a potential role in neuronal protein homeostasis and in turn, in neurodegeneration. To investigate the possibility, we performed a meta-analysis of RNA sequence data on neurodegenerative disorder models available in public databases. To identify appropriate groups of datasets, we performed methodical mining of RNAseq datasets in Gene Expression Omnibus (GEO) database as described in the methods. Fig. S3a is a bar chart showing the total number of RNAseq datasets in the GEO database with Neurodegeneration or Autophagy as the MeSH terms. As of March 2024, from humans and mice, respectively, 801 and 1023 neurodegenerative disorder and 262 and 225 autophagy datasets were available. Of the autophagy datasets, ∼10% (28 and 22 in humans and mice, respectively) have neurodegenerative diseases as a MeSH term, whereas <5% of the neurodegenerative datasets included autophagy as a MeSH term (Fig. S3b). Of the 28 human disease datasets, we chose 17, as described in the Methods, which we refer to as the ‘autophagy’ group of neurodegeneration datasets.

A second group of 15 neurodegeneration datasets were identified, which had no mention of autophagy in their metadata, as described in the Methods. These were referred to as the ‘non-autophagy’ group of neurodegeneration datasets. We hypothesized that the autophagy vs non-autophagy groups of neurodegeneration datasets were different owing to their disease etiology and progression. The two groups of datasets were subjected to data processing pipelines outlined in Fig S3c. A set of 90 genes consisting of 38 endo-lysosomal ion channels, 6 V-ATPase subunits and 46 autophagy related genes (ARGs) were selected for gene expression studies (Table S5). Table 2 shows relevant identifiers and metadata information for the autophagy and non-autophagy dataset groups. Fold changes (FC) in gene expression were measured in condition 2 with respect to condition 1.

**Table 2.** Metadata of datasets: Details of the chosen datasets in autophagy and non-autophagy groups, such as respective GSE ID, associated disease, RNA source, and two specific conditions with respective sample numbers in parenthesis for each condition. Abbreviated disease names are shown in parentheses in the disease column

| S.No | GSE ID | Disease | RNA Source | Condition 1<br>(Sample #) | Condition 2<br>(Sample #) |
| --- | --- | --- | --- | --- | --- |
| Autophagy Datasets |  |  |  |  |  |
| 1 | 239282 | Age-related Cognitive Disorders (ACD) | Patient Capillary Blood | Control Timepoint-1 (14) | ACD Timepoint-1 (16) |
| 2 | 160299 | Parkinson's (PD) | Patient Blood Plasma | Control (4) | PD (4) |
| 3 | 241430 | Age-related Huntington's (AHD) | Patient Fibroblasts induced Neurons | Young_MSN (6) | Old_MSN (6) |
| 4 | 194241 | Age related Huntington's (AHD) | Patient Fibroblasts induced Neurons | Young-MSNs (18) | HD-MSNs (18) |
| 5 | 208784 | GBA-Parkinson's (GBA-PD) | Patient iPSC induced Midbrain Organoids | Control (6) | PD (6) |
| 6 | 238013 | Alzheimer's (AD) | CVB iPSC induced Neurons | CVB_iN_WT (4) | CVB_iN_KO (4) |
| 7 |  |  | CVB iPSC induced Astrocytes | CVB_iAst_WT (4) | CVB_iAst_KO (4) |
| 8 | 213289 | DEGS1 mediated Sphingolipidosis (DEGS-SL) | HAP1 cell line | HAP1 WT (3) | HAP1 DEGS1 KO (3) |
| 9 | 135743 | Idiopathic Parkinson's (iPD) | Appendix Tissue from patients | Control (47) | PD (12) |
| 10 | 144254 | Alzheimer's (AD) | Gray Matter from inferior parietal lobule of postmortem brain | E2c (8) | E4c (22) |
| 11 |  |  |  | E2c (8) | E33 (12) |
| 12 | 139989 | Parkin-Hypoxia (P-H) | HeLa cell line | mCherry Normoxia (3) | Parkin Hypoxia (3) |
| 13 | 125239 | Idiopathic Parkinson's (iPD) | Patient Fibroblasts induced Neurons | Control_iN (8) | PD_iN (9) |
| 14 | 107323 | Wilson Disease (WD) | Hep G2 cell line | wt cells untreated (3) | ATP7B KO treated with Cu (3) |
| 15 | 99559 | p62 knockout mediated defects (p62-KO) | Neurepithelial Stem Cell differentiated Neurons | NES control 4d (3) | NES KO 4d (3) |
| 16 | 199194 | Niemann-Pick Type A (NP-A) | Patient Fibroblasts | NPA_untrt (3) | NPA_SMtrt (3) |
| 17 | 184520 | Iron accumulation mediated disorder (NBIA) | Patient Fibroblasts | NHDF (3) | WDR45 (3) |
| 18 | 133323 | Corticobasal Degeneration (CBD) | iPSC induced Neural Progenitor Cells | Short Repeat (3) | Intermediate Repeat (3) |
| 19 | 145069 | Glaucoma | Patient iPSC induced Retinal Ganglion cells | H3_correction (4) | H3_E50K (4) |
| Non-Autophagy Datasets |  |  |  |  |  |
| 1 | 52946 | Amyotrophic Lateral Sclerosis (ALS) | Whole Lumbar Spinal Cord from patients | Healthy (10) | ALS (10) |
| 2 | 61405 | Huntington's (HD) | Whole Peripheral Blood from patients | Control (8) | HD (11) |
| 3 | 66573 | Relapsing-Remitting Multiple Sclerosis (RRMS) | Whole Peripheral Blood from patients | Control (8) | RRMS (6) |
| 4 | 75852 | Ataxia-Telangiectasia (A-T) | Patient iPSC derived neural progenitor cells | neural progenitor control (3) | neural progenitor ataxia (3) |
| 5 | 77598 | MERTK based Multiple Sclerosis (MERTK-MS) | Purified Monocytes from patient blood | Control (3) | MS (5) |
| 6 | 86654 | Parkinson's (PD) | H9 hESCs differentiated ventral midbrain progenitors | DA-Low (7) | DA-High (8) |
| 7 | 95343 | Huntington's (HD) | Patient iPSC induced neural cells | Control (3) | HD (4) |
| 8 | 97100 | Huntington's (HD) | Patient iPSC induced brain microvascular endothelial cells | Control (4) | HD (8) |
| 9 | 98288 | Amyotrophic Lateral Sclerosis (ALS) | VCP WT/Mut iPSC induced Motor neurons | CTRL d35 (3) | MUT d35 (3) |
| 10 | 122736 | Cockayne Syndrome (CS) | Patient derived Fibroblasts | CSB WT (4) | CSB m/m (4) |
| 11 | 126155 | Familial Dysautonomia (FD) | Patient derived Fibroblasts | Untreated (6) | Kinetin (6) |
| 12 | 143570 | Ataxia with Oculomotor Apraxia 2 (AOA2) | Patient-derived Lymphoblastoid cells | Control (4) | AOA2 (4) |
| 13 | 157676 | Niemann-Pick Type C (NP-C) | Patient iNSC induced Brain organoids | WT (3) | NPC (3) |
| 14 | 162384 | TFAM Overexpression (TFAM-OE) | Lung A549 cells | Control (3) | TFAM (3) |
| 15 | 171342 | Tau Accumulation (Tau-Acc) | Neuroblastoma SHSY5Y cells | Tau-WT (3) | Tau-K174Q (3) |

### Intra-and Inter-Group Heterogeneity adds noise to the data

Heterogeneity in the different datasets may be attributed to the following: 1. the underlying variations in the etiology and progression of neurodegenerative disorders, 2. the associated diverse pathways of autophagy, and 3. different experimental models and/or tools adopted to study each of these disorders. We posited that the inherent heterogeneity in the neurodegeneration datasets would manifest in the carefully curated set of 90 genes which potentially govern autophagy (see Methods and Table S5).

First, differential expressions for these 90 genes were analyzed in both the groups of datasets. Fig 5a is a box plot showing the median values and the dispersion ranges for the fraction of genes which were differentially expressed (DEGs) in autophagy vs non-autophagy datasets. The median value of the fraction of DEGs in the autophagy group was significantly higher [Median, interquartile range (IQR); 0.33, 0.19-0.6) vs the non-autophagy group [median, IQR; 0.24, 0.18-0.3). Fig. 5b is a scatter plot showing the distribution of the fraction of DEGs in the ‘autophagy’ vs ‘non-autophagy’ dataset groups. In 6 of the datasets in the autophagy group (above dotted line) more than 50% of the genes were differentially expressed, while only 1 dataset from the non-autophagy group had greater than 50% of the genes differentially expressed. Fig 5c are the box plots showing the distributions of log_2_FC for all the 90 chosen genes in autophagy (left) vs non-autophagy (right) dataset groups. In the autophagy group, at least 20 out of the 90 genes showed >1.5-fold change (dotted line: log_2_FC = ±0.59) in as many as 9 out of 19 datasets. In contrast, in the non-Autophagy group only 2 out of 15 datasets showed >1.5-fold change in 20 out of the 90 genes.

**Fig. 5.**
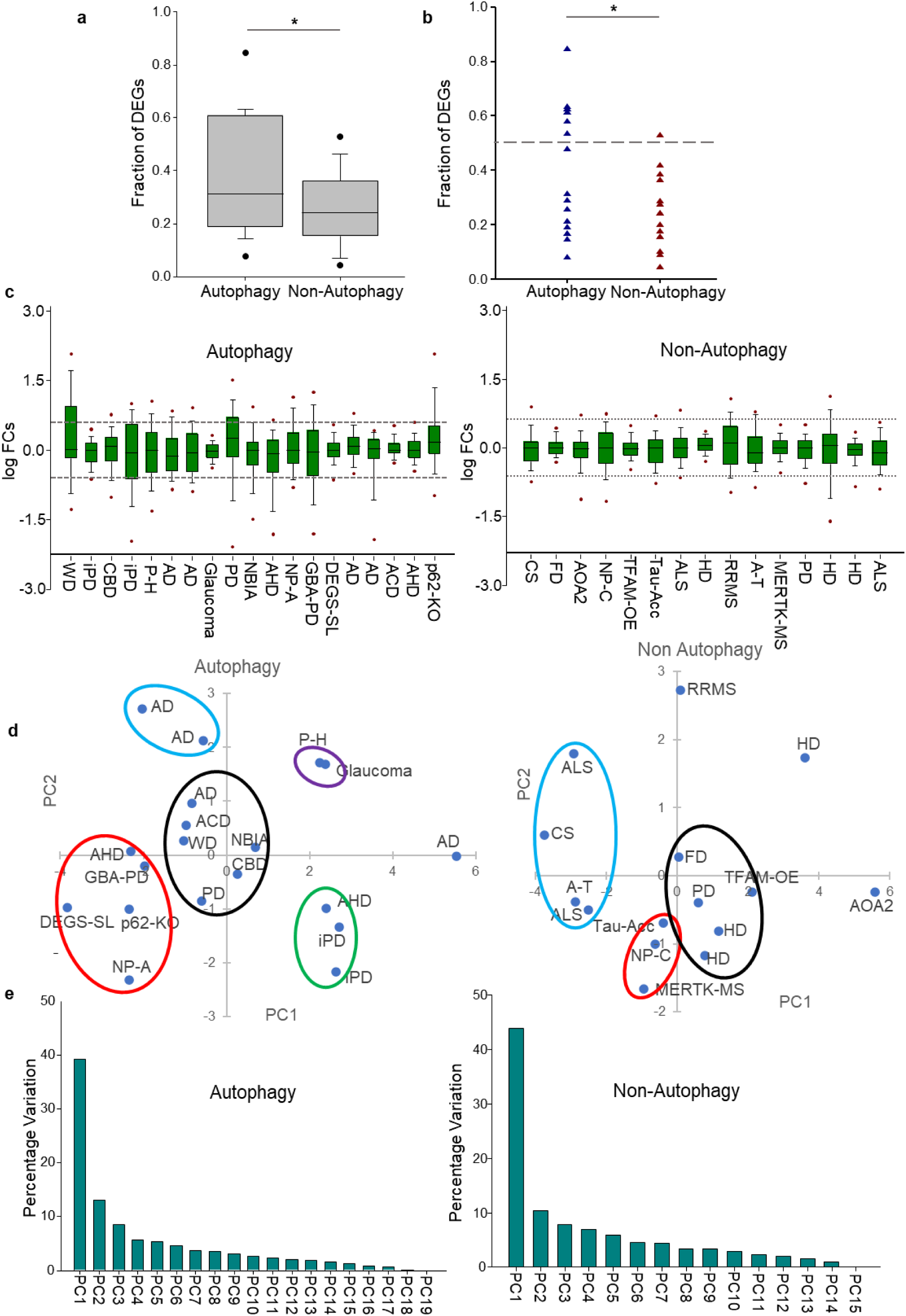
Heterogeneity in Neurodegeneration datasets: (a, b) Boxplot (a) and vertical scatter plot (b) showing the fraction of significant genes among the 90 chosen genes across all datasets in both autophagy and non-autophagy dataset groups. * = *p<0.05*. The dashed reference line in (b) corresponds to 0.5 fraction or 50% of the DEGs. **(c)** Boxplots showing the distribution of log fold changes of all the 90 genes in autophagy (left) vs non-autophagy (right) dataset groups. Red dots show the 5^th^ and 95^th^ percentile of each distribution. The dotted reference lines correspond to 1.5-fold change in either direction. **(d)** Scatterplot between principal components 1 (PC1) and PC2 showing similarity and heterogeneity between neurodegenerative datasets in the autophagy (left) vs non-autophagy (right) groups. Clusters were identified by k-mean algorithm and shown here using colored ellipses. **(e)** Bar plots showing the percentage variation explained by every principal component in autophagy (left) and non-autophagy (right) groups. *Abbreviations: ‘AD’ – Alzheimer’s Disease, ‘PD’ – Parkinson’s Disease, ‘iPD’ - Idiopathic Parkinson’s, ‘GBA-PD’ – GBA based Parkinsons, ‘HD’ – Huntington’s Disease, ‘AHD’ – Age-related Huntington’s, ‘ACD’ – Age-related Cognitive Disorders, ‘NP-A’ – Niemann Pick Type-A, ‘NP-C’ – Niemann Pick Type – C, ‘WD’ – Wilson’s Disease, ‘NBIA’ – Neurodegeneration with Brain Iron Accumulation, ‘CBD’ – Corticobasal Degeneration, ‘P-H’ - Parkin-Hypoxia, ‘DEGS-SL’ – DEGS1 based Sphingolipidosis, ‘p62-KO’ – p62 Knockout based defects, ‘CS’ – Cockayne Syndrome, ‘FD’ – Familial Dysautonomia, ‘AOA2’ – Ataxia with Oculomotor Apraxia 2, ‘TFAM-OE’ – TFAM Overexpression, ‘Tau-Acc’ - Tau Accumulation, ‘ALS’ – Amyotrophic Lateral Sclerosis, ‘RRMS’ – Relapsing-Remitting Multiple Sclerosis, ‘A-T’ – Ataxia-Telangiectasia, ‘MERTK-MS’ – MERTK based Multiple Sclerosis*

To understand the heterogeneity in the datasets, we performed principal component analysis (PCA) using log_2_FC of the 90 genes. Fig 5d (left and right) are scatterplots between the first two principal components (PCs), PC1 vs PC2 in both groups of datasets. To illustrate the heterogeneity in these datasets, the scatter data were clustered by using *k*-means algorithm (see Methods). The non-autophagy and the autophagy groups showed 3 vs 5 statistically significant clusters, respectively. The PC analysis resulted in meaningful clustering and segregation of datasets. For example, in the autophagy group, neurodegenerative disorders with strong genetic predisposition and/or due to genetic manipulation (for e.g. Age-related Huntington’s, GBA-Parkinson’s, DEGS1-sphingolipidosis, p62-knockout and Niemann-pick type A) were clustered together. Likewise, neurodegenerative disorders associated with abnormal storage of copper (Wilson’s disease, WD) and iron (Neurodegeneration with brain iron accumulation, NBIA) and are clustered together. On the other hand, different lineage of cells (astrocytes vs neurons) derived from embryonic stem cells in Alzheimer’s, and GBA-Parkinson’s (Parkinson’s due to anomaly in lysosomal acid beta-glucocerebrosidase) vs Parkinson’s induced by hypoxia are segregated widely (Fig. 5d).

In our analysis, PC1 and PC2 explained >50% of the variations whereas the top 5 PCs explained >70% of the variations in both the dataset groups (Fig 5e). Table S6 shows the top 5 contributing genes in both dataset groups to the top 5 principal components. Many of the top DEGs are important contributors to the principal components of both the dataset groups. In the autophagy group, the top PCs had higher number of ion channel genes including mucolipin family (MCOLN2/3) channels, voltage-gated Ca^2+^ (CACN), Cl^-^ (CLCN), inwardly rectifying K^+^ (KCNJ), cystic fibrosis transmembrane conductance regulator (CFTR), and transient receptor potential channels (TRPs) vs the non-autophagy group.

Fig S4a and S4b shows the top 12 ATGs/ARGs and ion channel genes, respectively, having >1.5-fold change in expression. However, these genes show marked heterogeneity in their levels of differential expression in the datasets of both the groups. Fig. S4a, S4b (Left) show that as many as 6 ATGs and all the 12 ion channel genes show differential expression in > 33% of the autophagy datasets. In contrast, only 1 ATG and 1 ion channel show differential expression in >33% of the non-autophagy datasets (Fig. S4a, S4b (Right)). In the autophagy vs non-autophagy groups, the following gene sets were observed to be differentially regulated: 1. ATGs including ATG16L2, MAP1LC3A, GABARAPL1, 2. Transcription factor TFEB and 3. Ion channel genes belonging to voltage-gated Ca^2+^, Na^+^, K^+^, volume regulated anion channels and transient receptor potential channels. Interestingly, the mucolipin family ion channel, MCOLN3 (TRPML3) was found to be differentially expressed in > 40% of all the neurodegenerative disorder datasets (including autophagy and non-autophagy datasets) (Fig. S4b). TFEB is the only transcription factor that was differentially regulated in >33% of the autophagy datasets.

### Gene Scoring and Correlation Networks

To identify the critical ATGs and ion channel genes across a heterogeneous group of datasets, we normalized the gene expression profiles for every gene in each dataset group (see Methods). Fig S4c shows the normalized scores of 90 genes in rank order for both groups. It shows that the normalized scores in the autophagy group are higher at every rank than the non-autophagy group. The difference in normalized scores (ΔS) = autophagy-non autophagy groups for every gene were calculated and shown in Fig S4d. For most of the genes, ΔS is positive, with CFTR showing the highest positive difference. Whereas TFEB showed the highest negative ΔS between autophagy vs non-autophagy groups. ATG5, SCNN1A, MCOLN2, and FOXO3 are the top genes that show high positive ΔS, whereas TFEB, LAMP1, KCNMA1, MCOLN3, and ATP6V0D1 (encodes for V-ATPase) are top genes that show high negative ΔS. The top 5 scoring ATG and ion channel genes in each category are shown in Table 3.

**Table 3.** Top 5 ATGs and ion channel genes with highest normalized scores in each dataset group.

| Rank | Autophagy |  | Non-Autophagy |  |
| --- | --- | --- | --- | --- |
|  | ATG/ARGs | Channels | ATG/ARGs | Channels |
| 1 | VAMP8 | CFTR | LAMP1 | TRPV6 |
| 2 | ULK1 | CACNA2D1 | MAP1LC3B | KCNMA1 |
| 3 | ATG5 | TRPV6 | TFEB | KCNJ8 |
| 4 | PRKAA2 | MCOLN2 | VAMP8 | MCOLN3 |
| 5 | STX17 | KCNJ8 | ATG9B | CACNA2D1 |

### Network Analysis to understand co-regulated gene expression

To identify the regulatory networks between these 90 genes, Pearson correlation coefficients (r) were determined between every pair of genes using their log_2_FCs. The coefficients were separately determined and shown for the autophagy and non-autophagy groups (see Fig S5a left and right). Regions of positive (e.g. 2^nd^ quadrant)/negative (e.g. 3^rd^ quadrant) correlation values appear scattered along the wings of the diagonal. To eliminate the possibility of these correlation coefficients arising due to random chance or background noise, the log_2_FCs in each dataset group were randomized. Fig S5b (left and right) shows correlation plots of the randomized log_2_FCs. We did not observe any significant correlation between any pair of genes in the randomized data. The correlation coefficients for some of the ATGs and channel genes are significantly higher in the autophagy vs non-autophagy groups. S5c is a subtraction matrix showing the difference in the *r* values between the autophagy vs non-autophagy groups.

Fig. S6a (left and right) shows correlation network plots for the 90 ATGs and channel genes in the autophagy vs non-autophagy groups using a correlation coefficient cutoff of *|r| > 0.55*. The ‘edges’ in the network correspond to the *r* - values for a pair of genes/‘nodes’. The total number of edges in the overall network were not significantly different between the autophagy vs non-autophagy groups. But the number of edges per nodes in the network between both the groups were significantly altered for many genes. S6b shows the overall changes in the number of edges for each gene between autophagy vs non-autophagy dataset groups in rank order. Of these, at least 37 genes had differences of more than 6 edges between the two groups (Fig 6a). It is important to note that a significant proportion of these genes are the top differentially expressed genes across all datasets.

**Fig. 6.**
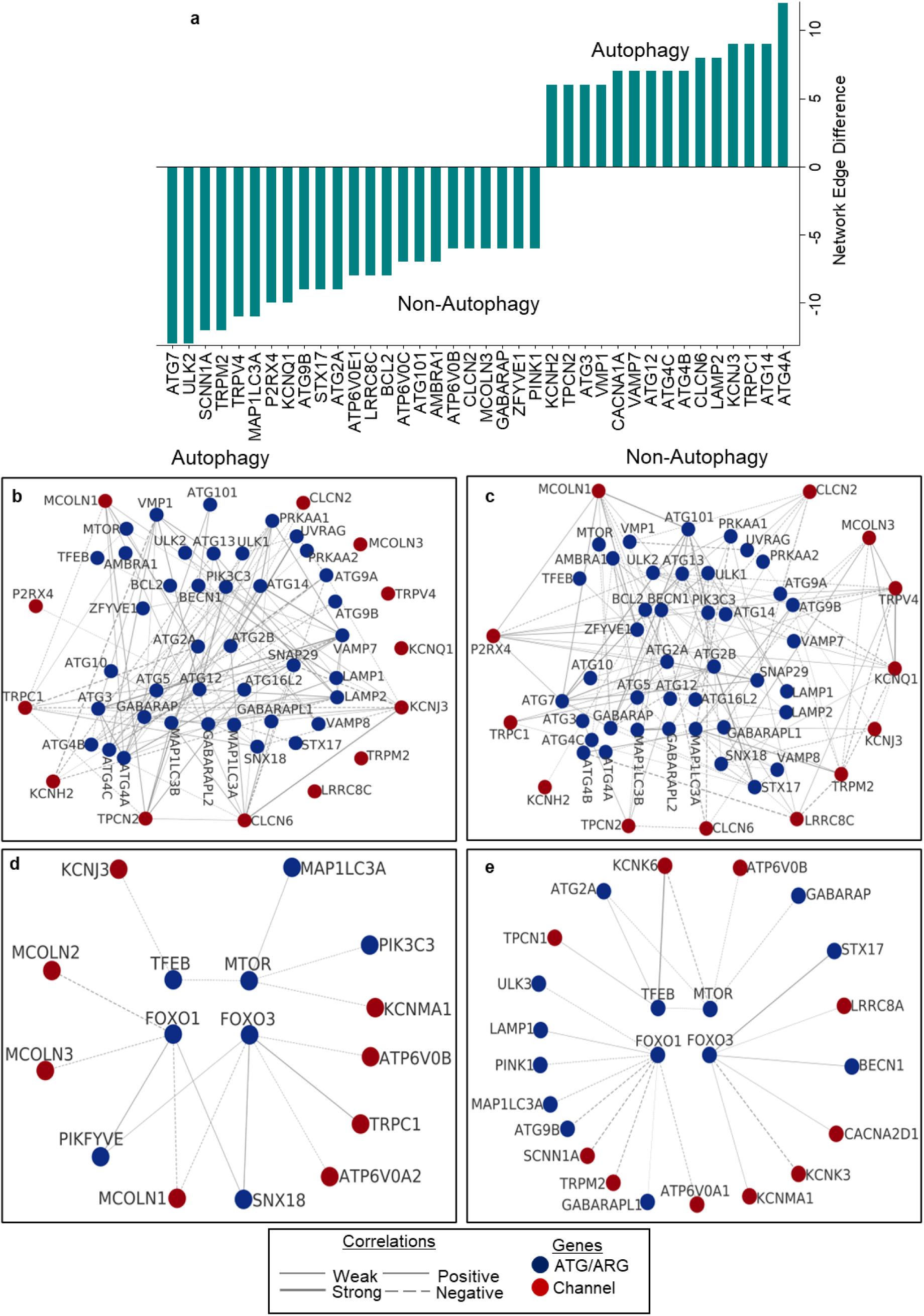
TRPML3 as a potential marker of familial neurodegenerative disorder: **(a)** Bar plot showing the top genes with large differences in number of network edges between autophagy and non-autophagy groups (autophagy minus non-autophagy). **(b, c)** Subnetworks of canonical autophagy pathway genes and selected endo-lysosomal ion channels from the fold change correlation networks of autophagy (d) and non-autophagy (c) dataset groups. Note the complete absence of ATG7 and lack of connections for MCOLN3 (TRPML3) in autophagy group. (**d, e**) Subnetworks of transcription factors (TFEB, FOXO1, FOXO3) and MTOR from the fold change correlation networks of autophagy (d) and non-autophagy (e) dataset groups. Note the reduction in the number of network edges for TFEB and FOXO1.

Mutations in canonical autophagy pathway genes have been strongly implicated in neurodegenerative diseases (58, 59). To understand the interplay between the canonical autophagy genes and ion channels, we constructed network clusters of these 2 sets of genes for autophagy vs non-autophagy groups (Fig. 6b,6c). Remarkably, ATG7, a core autophagy gene which was connected to >9 nodes (including ATG13, ATG101, ATG2A and ion channel MCOLN1) in the non-autophagy group, completely disappears in the cluster in the autophagy group. Likewise, ion channel genes MCOLN3 (TRPML3), TRPV4, KCNQ1, TRPM2, and LRRC8 which were connected to many other nodes in the network in the non-autophagy group completely lost their network in the autophagy group (Fig. 6b vs 6c). Further, transcription factors including TFEB (52–54), FOXO1 and FOXO3 (55) are known to regulate lysosome biogenesis, and promote autophagy induction. Therefore, we constructed network clusters for these transcription factors with ATGs and ion channel genes (Fig. 6d, 6e) for the autophagy vs non-autophagy groups. In the autophagy group the following network structures were observed: (1). TFEB was connected to fewer number of nodes in the autophagy group, (2). FOXO1 was negatively co-regulated with the mucolipin family of genes, MCOLN1, MCOLN2 and MCOLN3 in the autophagy group and (3) In the non-autophagy group, these transcription factor genes were co-regulated with denser network of ATGs and ion channels vs the autophagy group.

Further we identified ‘Hub’ genes using the DMNC algorithm (see Methods). They undergo significant alterations in network connectivity between the two dataset groups. Many of the top differentially expressed genes showed up as Hub genes. The top 15 Hub genes in both the networks, including crucial ATGs/ARGs, and endo-lysosomal ion channels are shown in Table 4.

**Table 4.** Top 15 hub genes identified using DMNC algorithm from both dataset groups.

| Rank | Autophagy | Non-Autophagy |
| --- | --- | --- |
| 1 | PIK3C3 | SNAP29 |
| 2 | KCNJ4 | ATP6V0D1 |
| 3 | ATG13 | ATG7 |
| 4 | KCNH2 | ATG13 |
| 5 | CACNA1A | GABARAP |
| 6 | BECN1 | LRRC8C |
| 7 | PACC1 | PRKAA1 |
| 8 | LAMP2 | PINK1 |
| 9 | ATG4C | ATG101 |
| 10 | CLCN7 | TMEM175 |
| 11 | GABARAPL1 | PIKFYVE |
| 12 | ATG12 | HVCN1 |
| 13 | VMP1 | MCOLN1 |
| 14 | ATG2B | MCOLN3 |
| 15 | SNAP29 | LAMP1 |

## DISCUSSION

The results presented here establish the role of TRPML3/MCOLN3 as a key autophagy regulator gene having direct or indirect roles in autophagy. The following lines of evidence establish the significance of TRPML3 in autophagy and/or neurodegeneration: (1) TRPML3 over-expression in a simple *in vitro* cell line model induced autophagy, and, more robustly, under serum starvation condition (Fig. 1). We observed TRPML3 localization and function in strategic cell compartments (Fig. 2) during autophagy induction similar to earlier observations by others (29, 60), (2) Ca^2+^ signals owing to the ion channel activity of TRPML3 was necessary for the induction of autophagy (Fig. 3), (3) TRPML3 over-expression in the HEK cells resulted in the enrichment of neuronal gene sets, including those responsible for axon guidance, synaptogenesis, sensory perception, neurotransmitter mechanism, and dendritic arborization (Fig 4), (4) TRPML3 induced TFEB mediated autophagy and altered the expression of neuron-specific transcription factors SOX2, ETV5, FOXC1 and 3, (5) In neurodegenerative disorders caused due to anomaly in the autophagy process, TRPML3 associated gene network was completely disrupted (Fig 6), and (6) TRPML3 was one of the key differentially expressed genes in genetic neurodegenerative disorders (14/20) irrespective of whether they are exacerbated by autophagy anomaly. DGE analysis shows that TRPML3 was the only channel gene to be differentially expressed in >40% of all the datasets (Fig S4). In a nutshell, TRPML3 seems to be a key genetic denominator in the familial neurodegenerative disorders that we have studied in this manuscript and, indirectly, plays a role in other sporadic or idiopathic neurodegenerative disorders owing to its role in autophagy.

The above conclusion is, however, based on only 32 datasets, which were selected from a pool of >1000 neurodegenerative disorder datasets. However, there was a strong rationale to ignore this large number of datasets for our analysis. Our data mining showed that only ∼5% of the neurodegenerative disorder conditions (RNA-seq datasets containing ‘autophagy’ as a MeSH term) could be directly linked to anomaly in autophagy processes (Fig. S3). This indicates that there is either a weak correlation between autophagy and neurodegeneration or a complex one which is not clearly understood. Autophagy reportedly plays conflicting roles (causative/protective) in neurodegenerative disorders (61). Our experimental design was driven to understand this dichotomy in the etiology of neurodegenerative disorders with respect to autophagy and hence, the datasets were carefully chosen and classified in two clear groups (Table 2).

We performed meta-analysis on the expression patterns for 90 genes (Table S5), with a focus on endo-lysosomal ion channels, transcription factors and ATGs to explore the global gene expression changes that underlie the 2 distinct classes of neurodegenerative disorders. The hypothesis was that if the classification was rational, the meta-analysis would be able to predict key driving genes (out of the above set) involved in neurodegenerative disorders. Our data mining showed that the autophagy group has more idiopathic/sporadic disorders (8/17) whereas most of the datasets in the non-autophagy group (12-14/15) were genetic in nature with known inheritance patterns (Table 2). This indicated that sporadic neurodegenerative disorders have a higher propensity of having impaired autophagy compared to hereditary disorders. Further, we report that the autophagy group has a higher fraction of DEGs, and larger fold change distributions for the selected 90 genes vs the non-autophagy group (Fig. 5). Meta-analysis is a powerful method. We have used a relaxed cutoff to extract significantly differentially expressed genes. It was necessary, but nevertheless a potential pitfall of the method, as adopted here. From the meta-analysis with the 90 genes, we have identified common patterns and could exhibit heterogeneity in autophagy across different disorders.

TFEB is the master regulator of autophagy. It is localized in the cytoplasm during conditions of nutrient abundance (62). Under starved condition TFEB gets dephosphorylated and shuttles to the nucleus. In the nucleus, it binds to the CLEAR motif of its target genes, many of which are involved in lysosomal biogenesis and autophagosome formation (53). Our results suggest that MCOLN3 and TFEB show large fold changes in the non-autophagy group, with TFEB being the largest among the 90 genes. We propose that TFEB is one of the key transcription factors that potentially drives the differential gene expression patterns in the autophagy vs non-autophagy groups. Gene regulatory networks reveal that ATG7, V-ATPase genes, and TRPML3 lost most correlations and TFs show significantly altered correlations in the autophagy network (Fig 6). Loss of ATG7 (initiator of autophagosome biogenesis) and V-ATPase correlations combined with suppressed TFEB regulation show that autophagosome biogenesis and lysosomal integrity are probably compromised leading to neurodegeneration in the autophagy group. For the non-autophagy group, we assume that autophagy is functional and behaves as a protective mechanism. ATG7 and V-ATPase genes, being the major hub genes in the non-autophagy network support that assumption (Fig. 6, Table 4). We propose TRPML3 as a genetic marker to differentiate neurodegenerative disorders with functional autophagy from disorders with impaired autophagy.

The *in vitro* serum-starvation model in HEK cells chosen for the current study was a suitable and clean system to study the role of TRPML3 in isolation due to the following reasons: (i) TRPML3, a *bona fide* endo-lysosomal ion channel, robustly expresses in the plasma membrane, (ii) The endogenous expression of TRPMLs in the HEK cells is low. Therefore, the observations and inferences drawn here about TRPML3 pertain mostly to the channel, (iii) Low pH is known to inactivate TRPML3 activity in the endo-lysosomal compartments. Therefore, TRPML3 expressed in the plasma membrane is a good system to eliminate this factor which might potentially occlude the inferences. However, we observed changes in the rectification properties of the TRPML3 currents during serum starvation (Fig. 3i). This indicated that either the surface expression of the channel was increased, or the biophysical properties of the channels were altered during serum starvation. Both possibilities suggest a potential role of post-translational modification of the channel during serum starvation. This needs to be investigated in the future. The bulk RNAseq experiments used to understand the mechanism of TRPML3 mediated gene expression in HEK293T cells is not an ideal method. Cellular heterogeneity owing to the variability in transfection and gene expression can potentially alter the degree of differential expression though the directionality of gene expression is less likely to change due to the robust statistical methods followed in this work. Serum starvation is known to induce autophagosome biogenesis via the established canonical pathway of autophagy, where autophagosome is formed at ER, undergoes trafficking through the endosomal system and finally fuses with lysosomes. Our microscopy studies show that a significant population of TRPML3 shuttles out of ER and ends in late endolysosomes upon starvation (Fig. 2) and there was a concomitant higher colocalization with LC3. These results are consistent with the canonical autophagy pathway.

Phosphoinositides are known to regulate TRPML family proteins. Activation of TRPML3 channels on plasma membrane under starvation cannot be conclusively ascribed to downregulation of the phosphoinositide PI(4,5)P_2_. While drosophila TRPML (63) and human TRPML1 (64) are reportedly blocked by PI(4,5)P_2_, there is no direct evidence of PI(4,5)P_2_ mediated blocking of TRPML3. Besides, PI(4,5)P_2_ (65, 66) and TRPML3 having functional commonality in regulating endocytosis and autophagy, makes it unlikely that PI(4,5)P_2_ blocks TRPML3 on plasma membrane.

From RNA-seq, we observed that serum-starvation induced autophagy downregulates development-related neuronal markers and TRPML3 overexpression significantly enhances the function of autophagy (Fig 4). Under fed conditions, TRPML3 overexpression upregulates development-related neuronal TFs thus exhibiting dual roles.

In summary, using meta-analysis we identified differences in regulation of autophagy between two distinct groups of neurodegenerative disorder datasets with different etiologies. We propose that TRPML3/MCOLN3 is a potential genetic marker for neurodegenerative disorders with functional autophagy. In an *in vitro* model, we show that TRPML3 promotes neuronal development under basal conditions while simultaneously arrests development by augmenting the function of starvation-induced autophagy. Overexpression and activation of TRPML3 directly induces autophagy while autophagy controls the localization and positively modulates the functional state of the channel. We propose that TRPML3 should be investigated as a potential marker for neurodevelopment and neurodegeneration.

## ACKNOWLEDGEMENTS

We are grateful to the Kim Lab, SKKU University, South Korea for generously gifting us the clones of TRPML3-EGFP, TRPML3-mCherry and LC3-EGFP. We are also thankful to the Menon lab, KSBS, IIT Delhi for sharing the pEGFP-C2 plasmid vector. We acknowledge the high-performance computing facility and the central research facility, IIT Delhi, respectively, for computational and experimental support. We thank Nucleome Informatics Pvt. Ltd., Hyderabad for support in RNA sequencing. The research was funded by SERB CRG (CRG/2022/007550), DBT (BT/PR47726/CMD/150/26/2023) and ICMR (IIRP-2023-0990) grant to TKN and DBT Ramaligaswami Re-entry Fellowship (BT/RLF/Re-entry/33/2019) to JS. SA and MM were generously supported by research fellowship from IIT Delhi.

## FOOTNOTE

### Data Submission

RNAseq data has been submitted to the Gene Expression Omnibus database (GEO Database: https://www.ncbi.nlm.nih.gov/geo/) with accession ID: GSE300275.

## METHODS

### Mining of public RNA-seq datasets

Using Gene Expression Omnibus’s (GEO) (67) advanced search builder (https://www.ncbi.nlm.nih.gov/gds/advanced), the total available RNA-seq datasets were queried using the following search filters: 1. *Dataset Type* – ‘Expression Profiling by High Throughput Sequencing’, 2. *Organism* – ‘Homo Sapiens’ OR ‘Mus musculus’ and 3. *MeSH* – ‘neurodegenerative diseases’ or ‘autophagy’. To further identify datasets with both MeSH terms, a search filter of ‘neurodegenerative diseases AND autophagy’ was applied, leaving other filters unchanged. These comprise the ‘autophagy’ group of neurodegenerative disorder datasets with autophagy as a MeSH term. A second group of datasets were identified using ‘neurodegenerative disease’ as *search by ontologies* term in the GEO RNA-seq Experiments Interactive Navigator (GREIN) database ((68), https://dev.ilincs.org/apps/grein/?gse=). A diverse group of neurodegenerative diseases and models were considered in this group of datasets. The term ‘autophagy’ was not present in any search field of these datasets. These comprise the ‘non-autophagy’ group of neurodegenerative disorder datasets where autophagy was not implicated. Single cell RNA-seq (scRNA-seq) datasets were omitted from our analysis to maintain uniformity in data normalization and analysis across bulk RNA-seq datasets. Datasets with less than 3 samples per condition were omitted to permit statistical testing.

### Public RNA-seq data processing pipeline

Raw Transcripts Per Million (TPM) count data for the datasets in both groups (autophagy vs non-autophagy) were downloaded from the GREIN database. Datasets with inaccessible count data were omitted. Lowly expressed genes were filtered and Trimmed Means of M-values (TMM) based normalization was performed using edgeR Bioconductor package in R (69–71). We estimated the normalized Counts Per Million (CPM) of the transcriptomes for each of the datasets. CPMs of a set of 94 genes, including autophagy-related genes (ATGs) and endolysosomal ion channel genes, were extracted in every dataset. 4 out of 94 genes were omitted as they had zero expression in more than two-thirds of the total datasets in one or both groups (autophagy vs non-autophagy). CPM fold changes (FCs) were then estimated for the remaining 90 genes in condition 2 (CPM2) with respect to condition 1 (CPM1) (FC = CPM2/CPM1, see Table 2) for every dataset. If a gene had zero expression in only one condition, CPM+1 was applied before FC estimation. If a gene was not expressed in a dataset, log2FC for the gene in that dataset was assigned as 0. Statistical significance was determined by the Mann-Whitney U (MWU) test using R. Cutoffs of 1.5-fold change (**|**log2FC**|** > 0.58) and MWU p-value < 0.1 were considered for Differential Gene Expression (DGE) analysis.

### Principal Component Analysis (PCA) and Cluster analysis

PCA was performed on the log2FCs of the above 90 genes. We performed PCA using the ‘PCA’ package from ‘sklearn’ python3 library. Percentage variation of each PC, and contribution of each of the 90 genes to all the PCs were extracted. The top 5 contributing genes to the top 5 PCs were identified (Table S6). Datasets were clustered based on PC1 and PC2 scores using the K-means clustering algorithm (72). The number of clusters were determined by the elbow method. It involved plotting the within-cluster sum of squares values vs the number of clusters (WCSS). The optimal value of *k* was identified as the point where rate of decrease in WCSS sharply changed (73).

### Normalization of gene expression

To normalize log fold changes of each gene across a hetereogeneous group of datasets, we adopted the following normalization method. First, log2FCs of all 90 genes were min-max normalized in every dataset between the values-1 to 1 (*n*logFCi,j for i^th^ gene in j^th^ dataset). Normalized log2FCs were then multiplied with negative logarithm of the associated MWU p-value (-log10pi,j for i^th^ gene in j^th^ dataset) to obtain individual scores for every gene in each dataset. The absolute values of these individual scores were summated and averaged with the number of datasets in which the gene exhibits non-zero expression (Ni for the i^th^ gene) to obtain the normalized score (*Si* for i^th^ gene) as follows:

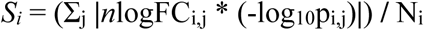

For the autophagy and non-autophagy dataset groups, normalized scores were calculated separately

### Correlation Analysis

Pearson correlation coefficients (PCCs) were determined between every possible pair of genes from their log2FCs using ‘numpy’ python3 library. These were referred to as the original coefficients. To estimate the effects of background noise, log2FCs were randomized 3 times and 3 sets of PCCs were determined using these randomized fold changes. They were then averaged to obtain the randomized coefficients. Original and randomized PCCs were determined separately for each dataset group.

### Network analysis

Fold change correlation networks between genes were constructed by applying a cutoff of **|**PCC**|** > 0.55 using Cytoscape software (74). Networks were constructed separately for each dataset group. The top 15 hub genes were identified using differential metabolic network construction (DMNC) algorithm (75) from the cytoHubba plugin (http://hub.iis.sinica.edu.tw/cytohubba) in Cytoscape. Specific subnetworks like canonical autophagy genes and transcription factors were manually extracted as required.

### Cell culture, Plasmids and Transfection

HEK293T cells were maintained in Dulbecco’s Modified Eagle’s Medium/Nutrient Mixture F-12 Ham (DMEM/F-12 Ham, Sigma Aldrich) supplemented with 1.2g/L NaHCO3 (Sigma Aldrich), 10% v/v Fetal Bovine Serum (FBS, Sigma Aldrich), and 1% v/v Penicillin-Streptomycin solution (Sigma Aldrich).

pEGFP-hTRPML3-C1, pEGFP-LC3-C1, and pCMV-TRPML3-mCherry-HA plasmids were generous gifts from Dr. Hyun-Jin Kim at Sungkyunkwan University (SKKU), South Korea. pEGFP-C2 plasmid was a generous gift from Prof. Manoj B. Menon at Indian Institute of Technology Delhi (IITD), India.

Transfection was performed using Effectene transfection reagent (Qiagen) according to manufacturer’s instructions for all the experiments described here except patch-clamp electrophysiology (see below). Cells were transfected at 40-50% confluency and harvested after 30-36 hrs. 700ng-2ug of DNA was transfected based on experimental requirements.

### RNA extraction and cDNA preparation

For RNA extraction, control vector/TRPML3 transfected HEK293T cells were seeded in 6-well plates. Approximately, 10^6^ cells were used to extract RNA for each of the replicates for different conditions. RNA was quantified by measuring absorbance at 260 nm (A260) and RNA purity was determined from the ratio of A260/A280. To induce starvation, cells were incubated in serum-free media for 4 hrs prior to RNA extraction. Four experimental conditions were generated: 10% serum fed and control vector overexpression (V), serum-starved and control vector overexpression (VS), 10% serum-fed and TRPML3 overexpression (T), serum-starved and TRPML3 overexpression (TS).

RNA was extracted using the RNeasy Plus and QIAshredder kits (Qiagen) as per the manufacturer’s protocols. From each sample, 1ug RNA was reverse transcribed to prepare cDNA using the iScript^TM^ cDNA Synthesis Kit (Bio-Rad). Reactions were performed in a total volume of 20ul. 4ul of 5X iScript Reaction Mix (containing primers and dNTPs) and 1ul of Reverse Transcriptase (RT) enzyme were used per reaction. Rest of the RNA was used for next-generation sequencing (Nucleome Informatics Pvt. Ltd., Hyderabad, India). cDNA was diluted in 1:2 or 1:3 ratios for qPCR experiments (see below).

### Experimental RNA-seq data processing, Differential Gene Expression (DGE) and Pathway Enrichment Analysis

RNA quality of the samples was assessed using agarose gel electrophoresis and the Bioanalyzer system (Agilent systems). Libraries were prepared by fragmenting the cDNA samples, specific adaptors were ligated and paired-end whole mRNA sequencing was performed on Illumina Novaseq 6000 platform. Triplicate samples corresponding to each of the 4 experimental conditions were sequenced.

The overall quality of the raw fastq sequence data was checked using FastQC (https://www.bioinformatics.babraham.ac.uk/projects/fastqc/) software. Adapters were trimmed using fastp tool (76). The quality of reads is measured by quality score. The quality score of a base is known as a Phred or Q score and is represented as Qphred=-10log10(e). “e” represents the sequence error rate and Qphred represents the base quality value. Reads were mapped to NCBI’s GRCh38 reference genome using HISAT2 tool (77). The alignment percentages for all fastq files were > 90%. Resulting.bam files were sorted, and transcript assembly was performed with Stringtie tool (78). Raw transcript counts were then extracted using prepDE.py function (78). The DESeq2 package in R was used to perform normalization, visualization and differential gene expression (DGE) analysis of the sample read counts (79). An alternate pipeline was also implemented where featureCounts tool (80) was used to estimate raw gene counts from HISAT2’s sorted.bam files. These counts were then subjected to DESeq2 based normalization and DGE analysis as before.

Differentially Expressed Genes (DEGs) were identified from both pipelines using the following cutoffs: p < 0.05 and adjusted p < 0.1 for Fed vs Starvation comparisons (‘V vs VS’ and ‘T vs TS’), p < 0.05 for Control vs Overexpression comparisons (‘V vs T and VS vs TS’). Among them, neuronal DEGs were identified based on a compiled gene list using specific Gene Ontology (GO) terms (81) and manual curation. The GO terms ‘generation of neurons’, ‘nervous system development’, ‘neurogenesis’, and ‘neuron differentiation’ were used for neuronal gene compilation. The gene lists from these GO terms were extracted using AmiGO 2 browser (https://amigo.geneontology.org/amigo) Manually curated genes were selected based on the description of their neuronal roles in NCBI and GeneCards (82) databases.

Pathway enrichment analysis on DESeq2 normalized counts from both pipelines was performed using Gene Set Enrichment Analysis (GSEA) tool (83). All gene sets under C2 – curated gene sets (7411 gene sets curated from various sources, including online pathway databases and the biomedical literature) and C5 – ontology gene sets human collections (Gene sets that contain genes annotated by the same ontology term) hosted by Molecular Signatures Database (MSigDB) (84) were assessed for enrichment. Enriched gene sets were filtered with a cutoff of p < 0.05. Neuronal genesets were identified manually.

### Realtime qPCR

Realtime quantitative PCR gene expression quantifications were performed and reported as per the MIQE guidelines (85). In all the experimental and control groups (V, VS, T, TS), RT-qPCR reactions were performed in a total volume of 10μl, comprising of 5µl 2X iTaq^TM^ Universal SYBR Green Supermix (Bio-Rad), 10ng cDNA template and 400nM of each primer (final concentration) on the CFX Opus Real-Time PCR platform (Bio-Rad). For each biological replicate, RT-qPCR reactions on the target genes were performed in technical triplicates. Gene expression levels were normalized to the average expression of GAPDH and b-Actin housekeeping genes. ΔΔCt method was utilized to determine fold changes between experimental conditions (86). *Outliers in ΔΔCt values were identified by Grubbs test and removed from final analysis.* The list of gene targets, their respective primers and size of the amplicons are given in Table S7. Primers were designed using Primer3 software (https://primer3.ut.ee/)

### Cell cycle analysis

HEK293T cells were seeded in 6-well plates for cell-cycle analysis. For serum-starvation, normal media was replaced with serum-free media at 4hrs prior to sample preparation. Cells were pelleted and then fixed with 70% prechilled Ethanol. Cells were then washed and resuspended in Dulbecco’s Phosphate Buffered Saline (DPBS, Sigma-Aldrich), followed by RNAse A (HiMedia) treatment. Finally, they were stained with Propidium Iodide (HiMedia) before sorting the cells in the BD LSRFortessa^TM^ X20 Flow cytometer (BD Biosciences). For nocodazole treatment conditions, cells were treated with either 75ng/ml or 100ng/ml concentrations (45) of nocodazole (Sigma Aldrich) for 48hrs. The proportions of cells in G1-S-G2/M phases were determined using the BD FACSDiva^TM^ software (BD Biosciences, San Jose, CA).

### Confocal Microscopy, Immunocytochemistry, and Image Analysis

HEK293T cells transiently transfected with LC3/TRPML3 were seeded onto glass coverslips coated with poly-D-lysine (PDL, Gibco) for microscopy experiments. Cells under different experimental conditions were fixed with 4% (v/v) paraformaldehyde (Sigma-Aldrich) in DPBS for 30min. For experiments involving Lysotracker, cells were loaded with 100nM Lysotracker Green DND-26 (Thermo-Fisher) for 90 min prior to fixation. For immunocytochemistry, cells were permeabilized with 0.05% Triton X-100 (Sigma-Aldrich) in DPBS for 5min. Cells were blocked with 5% Bovine Serum Albumin (BSA, HiMedia) suspended in 0.05% Triton X-100 in DPBS. Primary antibody incubation was done overnight at 4^0^C followed by fluorescent secondary antibody incubation (1:1000) at room temperature for 1hr. For nuclei staining, cells were loaded with 500ng/ml DAPI (Sigma-Aldrich) for 10min. Coverslips were then mounted onto glass sides with Mowiol 4-88 (Sigma-Aldrich) as a mounting medium. The following primary antibodies were used for ICC experiments: anti-TRPML3 (1:200, ACC-083, Thermo-Fisher), anti-KDEL-AF488 (1:300, PA1-013-A488, Thermo-Fisher), anti-ATG7 (1:200, ZRB1235, Sigma-Aldrich), and anti-LC3A/B (1:500, PA5-115500, Thermo-Fisher)

For colocalization studies, confocal images were captured using Leica TCS SP8 microscope with a 63X oil-immersion objective and 2x digital zoom. For puncta analysis, images were captured using ImageXpress Pico (Molecular Devices) with a 40X objective. Raw Z-stacked images were processed and analyzed using Fiji software (87). Preprocessing involved maximum intensity projection of z-stacks followed by background correction. Manders coefficients (88) were used as indicators for colocalization studies using JACoP plugin ((89), https://imagej.net/plugins/jacop). Puncta analysis was performed using ‘Find Maxima’ option in Fiji software. Cell counting was performed using ‘Analyze Particles’ option on DAPI filter.

### Calcium Assays

Control or transfected HEK293T cells were seeded at a density of 3*10^4^ per well onto PDL coated, clear bottom 96-well black plates (Corning-3603) for calcium assay experiments. Compounds of desired concentrations were prepared in a separate 96-well plate. Cells were incubated in Calcium 5 Assay dye (Molecular Devices) for 1hr. For EGTA-containing experiments, cells were incubated in Calcium dye with EGTA and devoid of calcium and magnesium salts. Calcium recordings were then performed using FLIPR-Penta High-Throughput Cellular Screening System (Molecular Devices). Camera gain of 2 and’Normal’ acquisition mode of the HS-EMCCD camera were applied during recordings.

### Whole cell Patch-clamp Electrophysiology

HEK293T cells were seeded onto PDL coated glass coverslips for patch-clamp experiments. Transfection was performed using Calcium Phosphate method (90). Patch-pipettes of 5-7 MΩ resistance were pulled from thick-walled borosilicate capillaries (Sutter Instrument) using a P-30 vertical micropipette puller (Sutter Instrument) by a two-stage pulling protocol. Patch pipettes were filled with the solution: 140mM CsCl, 2mM MgCl2, 10mM HEPES and 10mM EGTA and cells were placed in a bath solution comprising of 140mM NaCl, 5mM KCl, 10mM HEPES, 2mM MgCl2, and 2mM CaCl2 (7.4 pH adjusted with NaOH). Current recordings were obtained in voltage-clamp mode using dPatch amplifier (Sutter Instrument). Data acquisition and analysis was performed using the SutterPatch software. Regression analysis was performed using SigmaPlot 10.0.

### Statistical Tests

For public RNA-seq meta-analysis, MWU test was used as detailed before. For experimental RNA-seq data, statistical testing was performed by the DESeq2 package in R. For the rest of the data, Welch’s t-test was used for determining statistical significance.

