## Supplementary Figures and Tables for "TRPML3 regulates neuronal gene expression in an *in vitro* model of autophagy and may act as a genetic marker of familial neurodegenerative disorders"

**a**

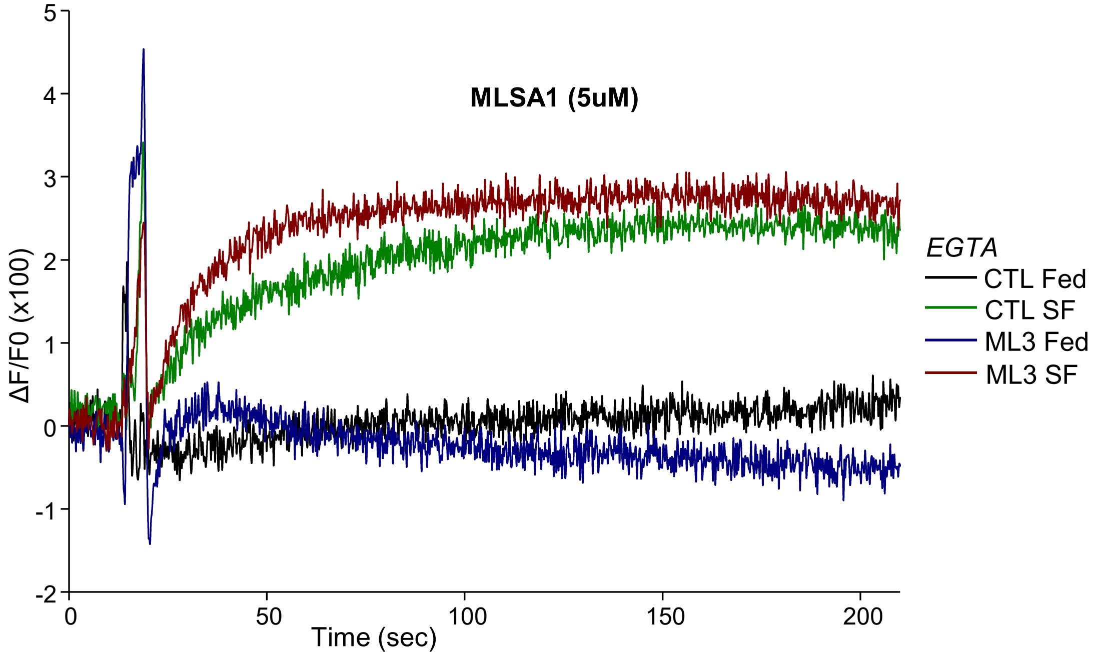


**Fig S1. Calcium Assays: (a)** Representative traces of calcium release from HEK293T cells under labelled experimental conditions upon treatment with 5uM of TRPML agonist MLSA1. Cells were incubated in dye devoid of $Ca^{+2}/Mg^{+2}$ ions and supplemented with EGTA. *Data was acquired of 6 wells of cells for each experimental condition*

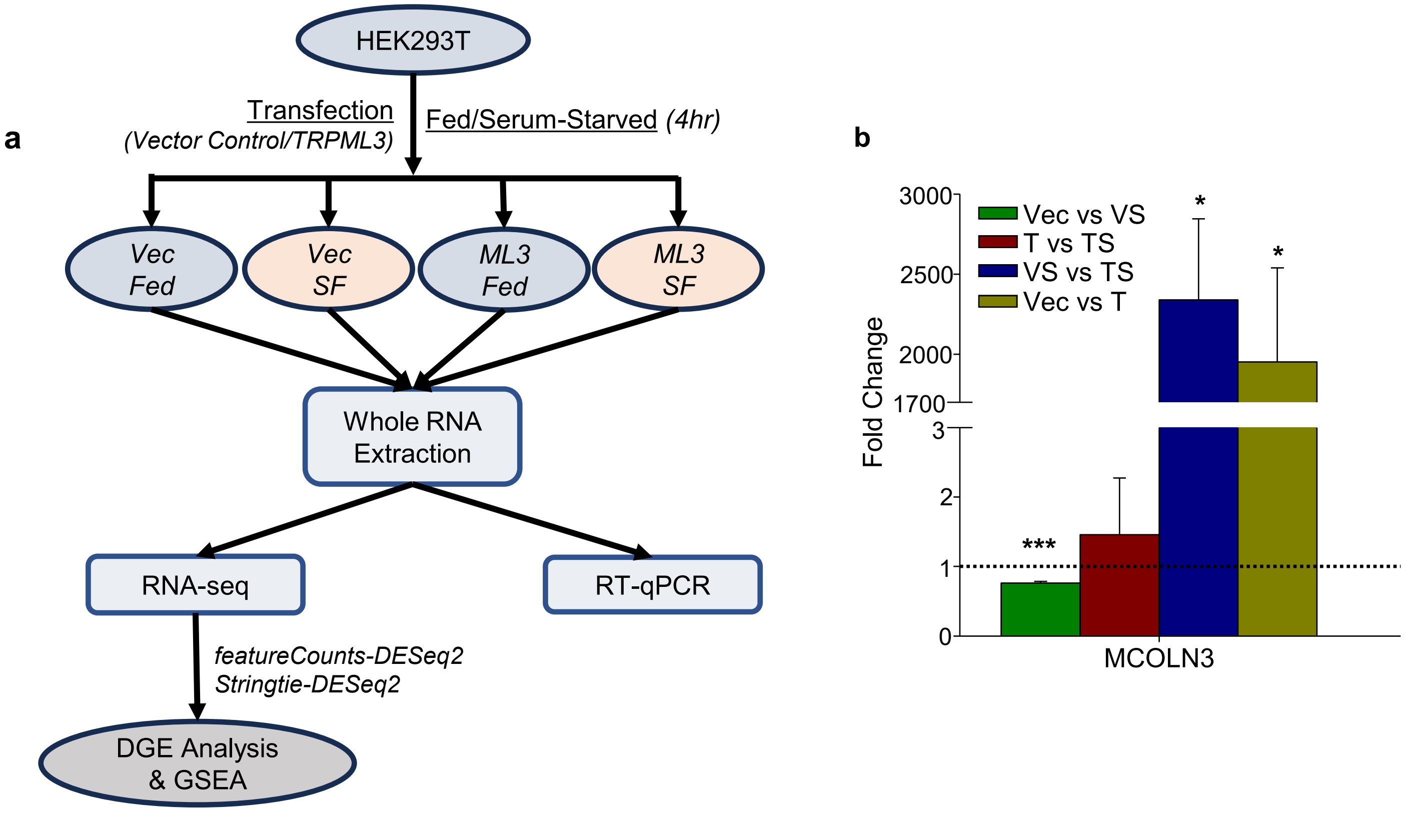


**Fig S2. MCOLN3 Overexpression Validation: (a).** Flowchart detailing the experimental approach for investigating the *in-vitro* serum-starvation model of autophagy. **(b)** qPCR validation of MCOLN3/TRPML3 overexpression in the *in vitro* serum-starvation model of autophagy. *** - $p < 0.005$, * - $0.01 < p < 0.05$

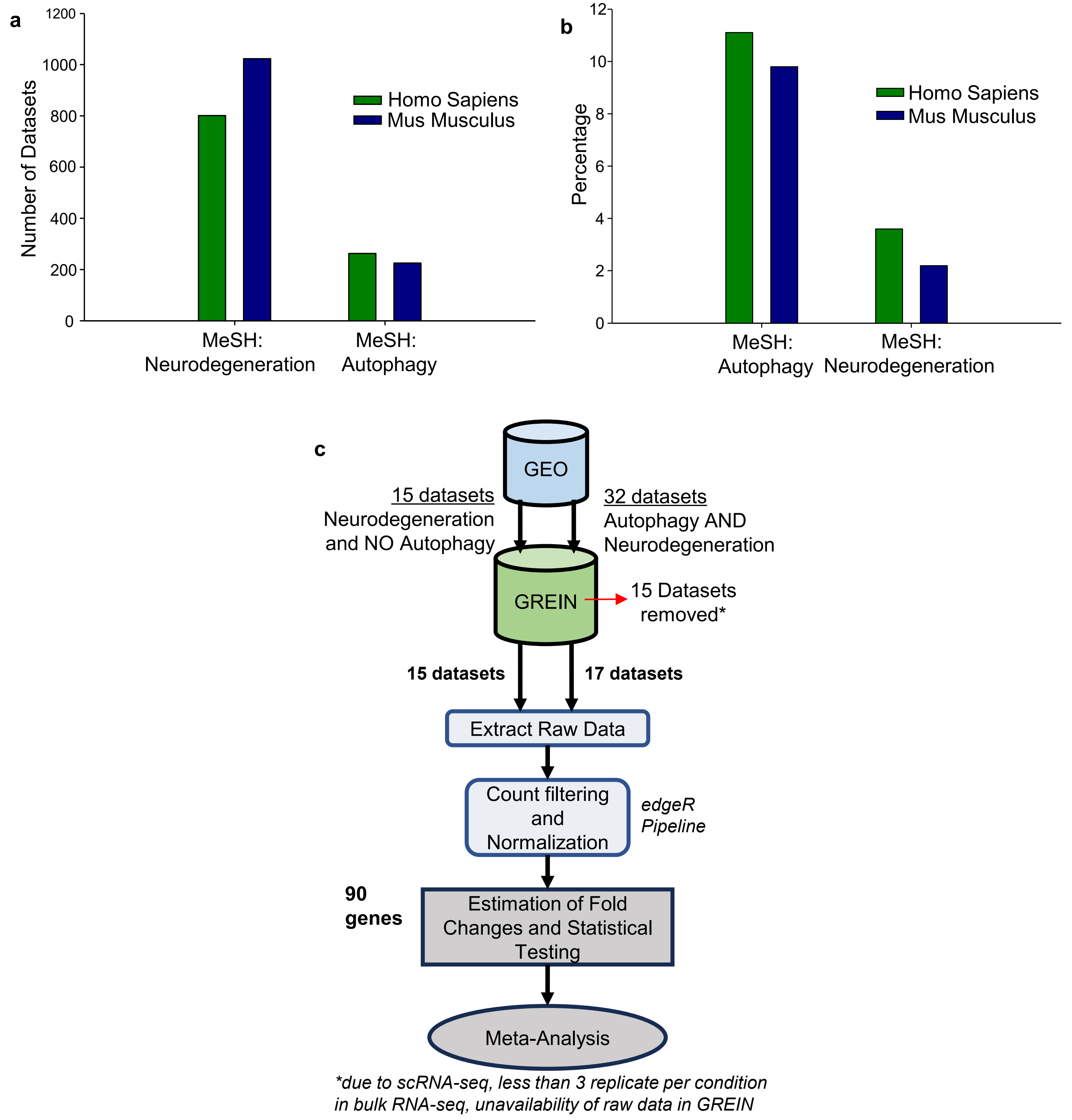


**Fig S3. Data Mining and Processing:** **(a)** Barplot showing the total number of RNA-seq datasets available in GEO with respective MeSH terms and organisms. **(b)** Barplot that shows the percentage of datasets with BOTH MeSH terms amongst total datasets from (a) with individual MeSH terms. **(c)** Flowchart detailing data acquisition and processing pipeline for meta-analysis. 90 genes (46 ATGs, 38 endo-lysosomal channel genes and 6 V-ATPase subunit genes) were chosen for downstream analyses.

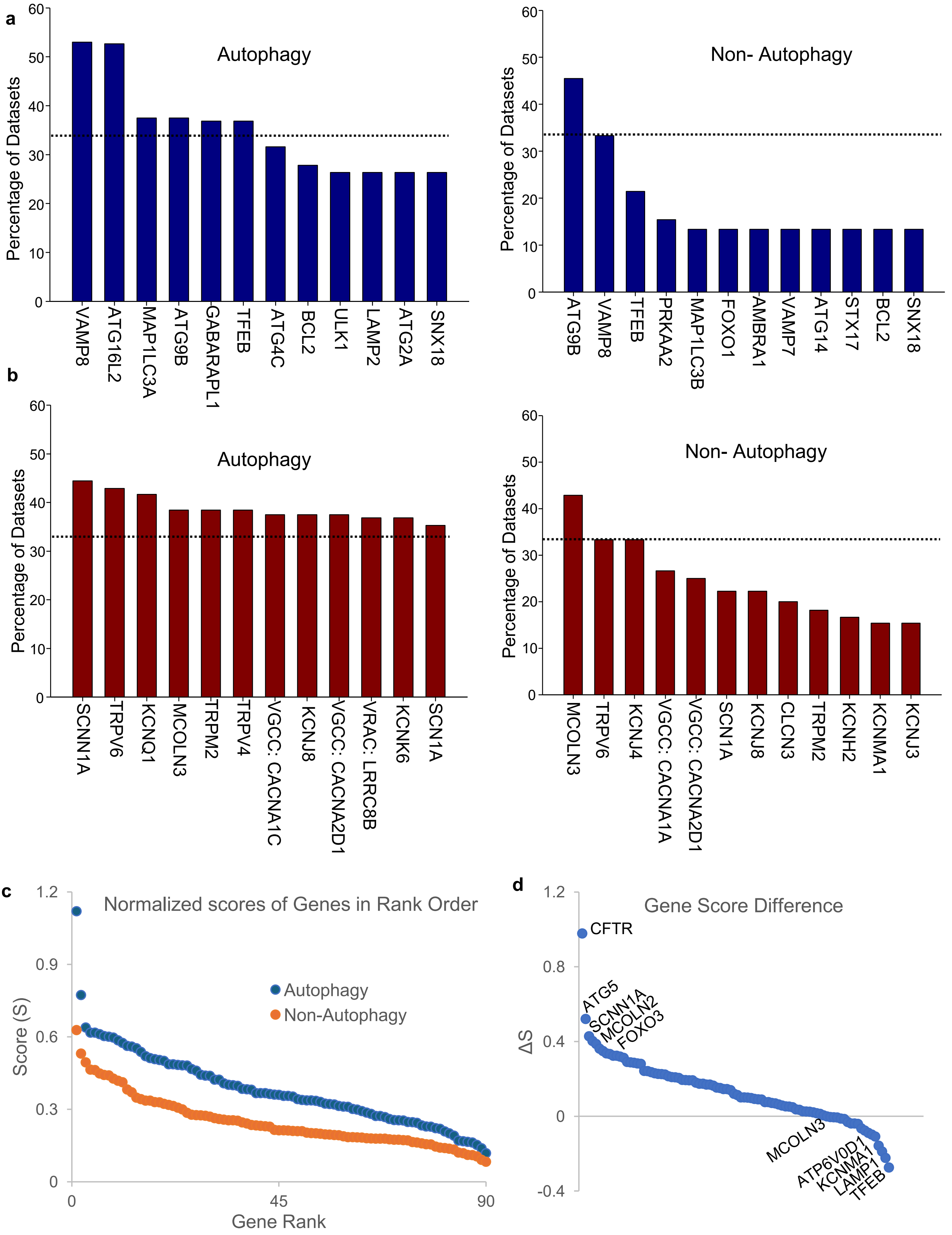

a
Autophagy
Percentage of Datasets
VAMP8
ATG16L2
MAP1LC3A
ATG9B
GABARAPL1
TFEB
ATG4C
BCL2
ULK1
LAMP2
ATG2A
SNX18
Non- Autophagy
Percentage of Datasets
ATG9B
VAMP8
TFEB
PRKAA2
MAP1LC3B
FOXO1
AMBRA1
VAMP7
ATG14
STX17
BCL2
SNX18
b
Autophagy
Percentage of Datasets
SCNN1A
TRPV6
KCNQ1
MCOLN3
TRPM2
TRPV4
VGCC: CACNA1C
KCNJ8
VGCC: CACNA2D1
VRAC: LRRC8B
KCNK6
SCN1A
Non- Autophagy
Percentage of Datasets
MCOLN3
TRPV6
KCNJ4
VGCC: CACNA1A
VGCC: CACNA2D1
SCN1A
KCNJ8
CLCN3
TRPM2
KCNH2
KCNMA1
KCNJ3
c
Normalized scores of Genes in Rank Order
Score (S)
Autophagy
Non-Autophagy
Gene Rank
d
Gene Score Difference
ΔS
CFTR
ATG5
SCNN1A
MCOLN2
FOXO3
MCOLN3
ATP6V0D1
KCNMA1
LAMP1
TFEB

**Fig S4. Top DEGs and Gene Scoring:** **(a)** Barplots showing the percentage of datasets for top 12 ATGs where they are differentially regulated with at least 1.5-fold change amongst the Autophagy (left) and Non-Autophagy (right) groups. **(b)** Barplots showing the percentage of datasets for top 12 channel genes where they are differentially regulated with at least 1.5-fold change amongst the Autophagy (left) and Non-Autophagy (right) groups. Dotted reference lines represent 33% of datasets. VGCC – Voltage gated Calcium Channel. VRAC – Volume Regulated Anion Channel. **(c)** Scatterplot shows the scores in rank order for all 90 genes in both Autophagy and Non-Autophagy dataset groups. Determination of scores is detailed in materials and methods section. **(d)** Scatterplot that shows the score difference (Autophagy minus Non-Autophagy) for each gene in decreasing order. Top 5 highest and top 4 lowest ranked genes along with MCOLN3 are labelled.

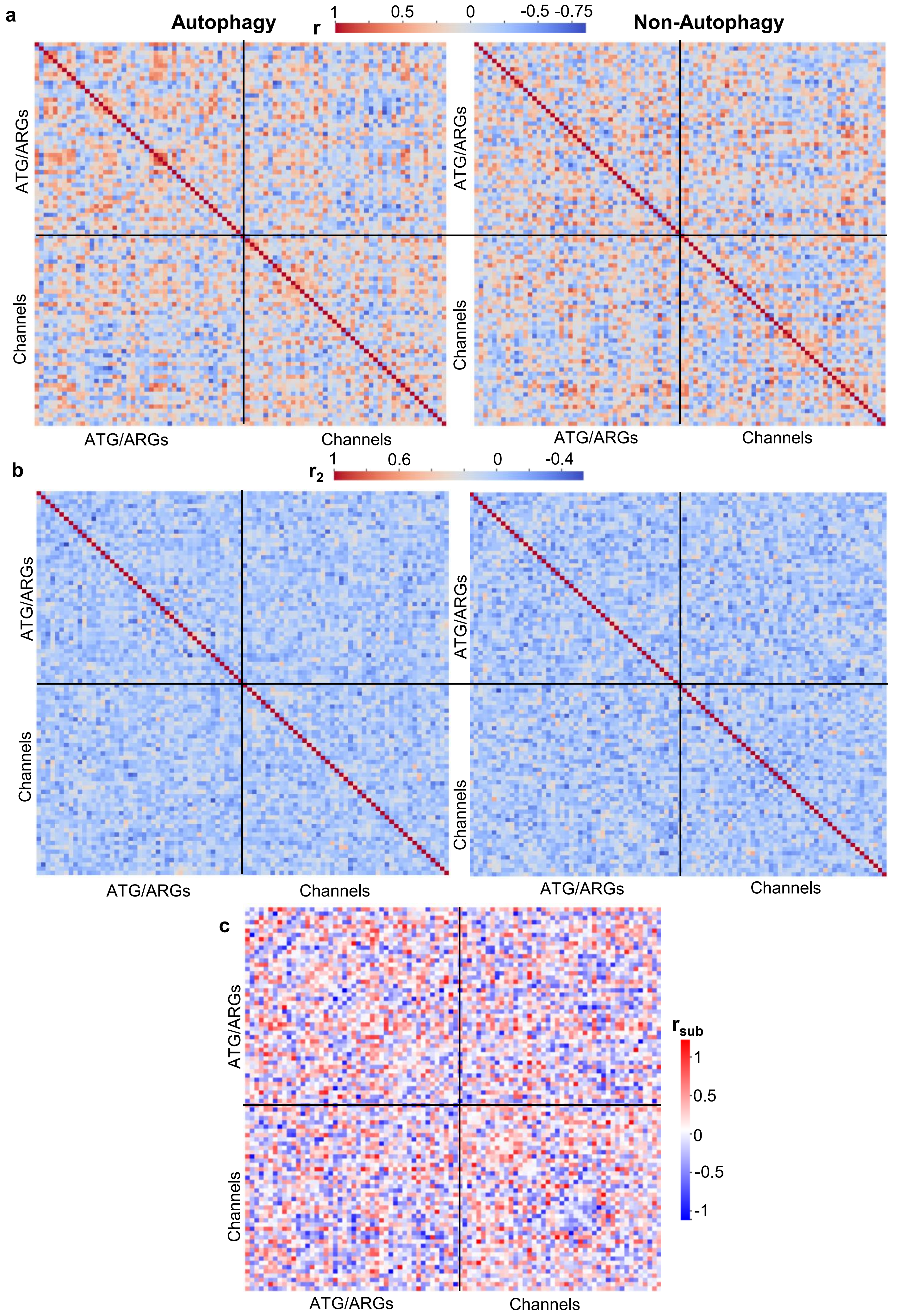

a
Autophagy
r
1
0.5
0
-0.5 -0.75
Non-Autophagy
ATG/ARGs
Channels
ATG/ARGs
Channels
ATG/ARGs
Channels
ATG/ARGs
Channels
b
$r_2$
1
0.6
0
-0.4
ATG/ARGs
Channels
ATG/ARGs
Channels
ATG/ARGs
Channels
ATG/ARGs
Channels
c
ATG/ARGs
Channels
ATG/ARGs
Channels
$r_{sub}$
1
0.5
0
-0.5
-1

**Fig S5. Correlation Analysis: (a)** Heatmaps of the original Pearson correlation coefficients (r) determined using log fold changes of every possible pair of genes in the autophagy (left) and non-autophagy (right) group of datasets. **(b)** Heatmaps of the randomized Pearson correlation coefficients ($r_2$) determined using log fold changes of every possible pair of genes in the autophagy (left) and non-autophagy (right) group of datasets. For all heatmaps, the halves on either side of the autocorrelation lines are mirror images. **(c)** Heatmap of difference in r values ($r_{sub}$) between autophagy and non-autophagy dataset groups.

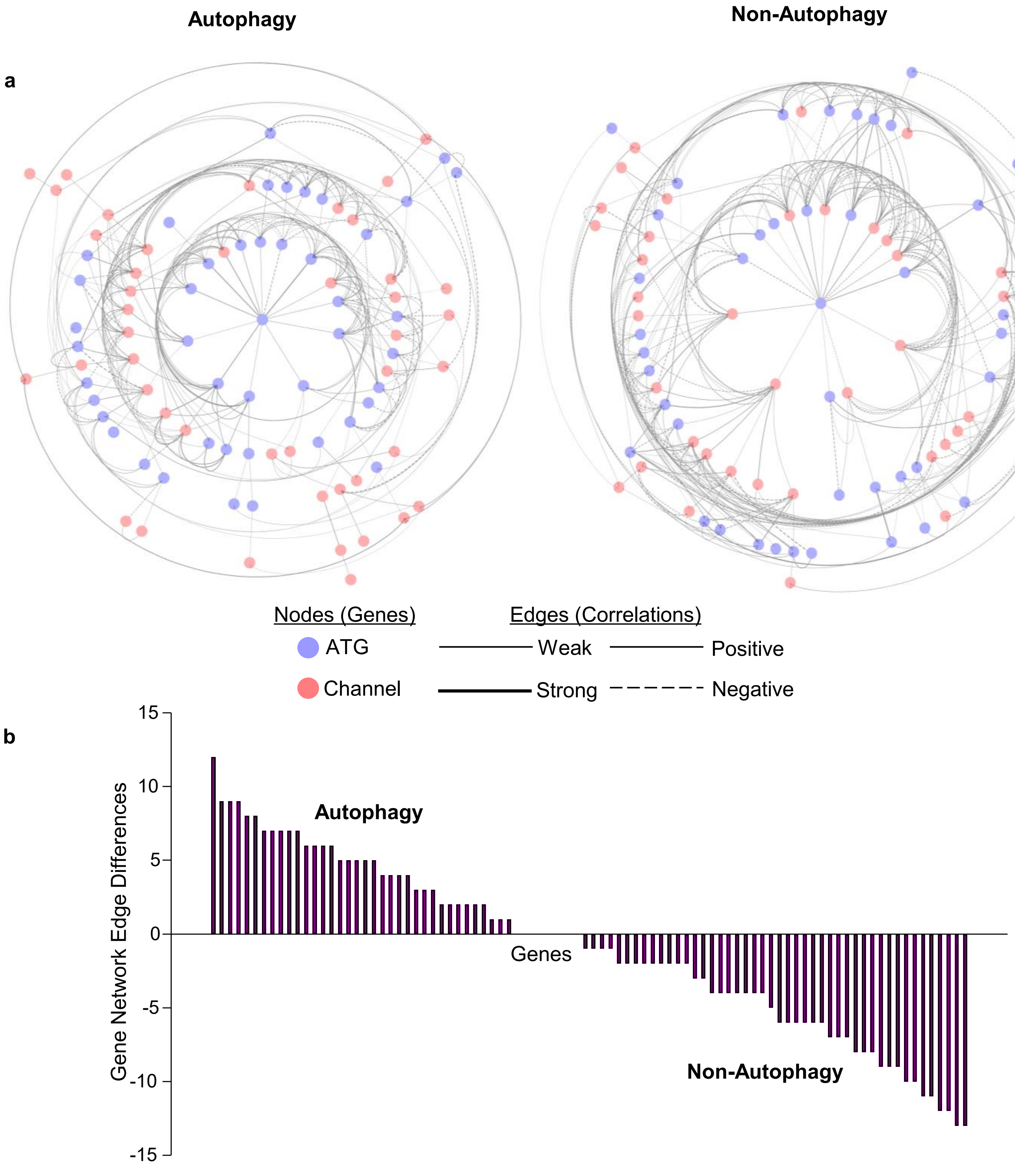


**Fig S6. Network Analysis: (a)** Overall fold change correlation networks constructed from PCCs with an applied cutoff of |PCC| > 0.55 in the Autophagy (left) and Non-autophagy (right) group of datasets. **(b)** Barplot representing the difference in number of edges (correlations) of all 90 genes in decreasing order between correlation networks of Autophagy vs Non-autophagy dataset groups (Autophagy minus Non-autophagy). *PCC – Pearson Correlation Coefficient*

| **Neuronal Genes** | **V vs VS** | **T vs TS** | **V vs T** | **VS vs TS** |
|---|---|---|---|---|
| Transcription Factors | | | | |
| ETV1 | | ↓ | ↑ | |
| ETV5 | | ↓ | ↑ | ↑ |
| ZNF365 | | ↓ | | |
| GDF7 | | ↓ | | |
| NR4A3 | | ↓ | | |
| DIXDC1 | | ↓ | ↑ | |
| SOX2 | | ↑ | | |
| FOXC1 | ↑ | ↑ | | |
| Structural and Signaling Proteins | | | | |
| NAB2 | | ↓ | ↑ | ↑ |
| RAPGEFL1 | | ↑ | | |
| LRRTM2 | | ↑ | | |
| MARK2 | | ↑ | | |
| LIF | | | ↓ | ↓ |
| NINJ2 | | | ↑ | |
| COL6A5 | | ↑ | ↑ | ↑ |
| ANKRD24 | | | ↑ | |
| Mutations leading to Neurological disorders | | | | |
| C9orf72 | ↓ | ↓ | | |
| FGD4 | | ↓ | | |
| CLIC2 | | ↑ | ↓ | |
| ZNF699 | | ↓ | | |
| SETX | | ↓ | | |

**Table S1.** List of selected neuronal DEGs with respective expression patterns in 4 pairwise comparisons. They are grouped into 3 categories. Red upward arrows correspond to significant upregulation while blue downward arrows correspond to significant downregulation in that specific comparison.

| C2_Geneset | T vs TS | V vs VS | V vs T | VS vs TS |
|---|---|---|---|---|
| FLORIO_HUMAN_NEOCORTEX | T | | | |
| FLORIO_NEOCORTEX_BASAL_RADIAL_GLIA_UP | T | | | |
| KEGG_AXON_GUIDANCE | T | | | |
| KEGG_NEUROACTIVE_LIGAND_RECEPTOR_INTERACTION | T | V | | |
| LEE_NEURAL_CREST_STEM_CELL_DN | | V | | |
| LEIN_CEREBELLUM_MARKERS | | V | | |
| LEIN_CHOROID_PLEXUS_MARKERS | | V | | |
| REACTOME_GABA_B_RECEPTOR_ACTIVATION | T | V | | |
| REACTOME_GABA_RECEPTOR_ACTIVATION | T | V | | |
| REACTOME_LONG_TERM_POTENTIATION | T | | | VS |
| REACTOME_NEUROTRANSMITTER_RELEASE_CYCLE | T | | | |
| REACTOME_SENSORY_PERCEPTION | | | | VS |
| WP_CELL_LINEAGE_MAP_FOR_NEURONAL_DIFFERENTIATION | T | | | |
| WP_DOPAMINERGIC_NEUROGENESIS | T | | | |
| WP_INTRACELLULAR_TRAFFICKING_PROTEINS_INVOLVED_IN_CMT_NEUROPATHY | T | | | |
| WP_NEURAL_CREST_CELL_MIGRATION_DURING_DEVELOPMENT | T | | | |
| WP_NEURAL_CREST_DIFFERENTIATION | T | | | |
| WP_NEUROINFLAMMATION_AND_GLUTAMATERGIC_SIGNALING | T | | | |
| WP_PARKINSON_39_S_DISEASE_PATHWAY | | | T | |
| WP_PHOSPHODIESTERASES_IN_NEURONAL_FUNCTION | T | | | |
| WP_SPINAL_CORD_INJURY | T | | | |

**Table S2.** List of enriched neuronal genesets from the ‘C2’ group (KEGG, REACTOME, and others) of curated genesets from Molecular Signatures Database (MSigDB) obtained from GSEA. The denoted condition under each pairwise comparison for each geneset represents the enriched condition. They are called as significant by both RNA-seq pipelines.

| C5_Geneset | T vs TS | V vs VS | V vs T | VS vs TS |
|---|---|---|---|---|
| GOBP_AUTONOMIC_NERVOUS_SYSTEM_DEVELOPMENT | T | | | |
| GOBP_AXON_GUIDANCE | T | | | |
| GOBP_DENDRITE_EXTENSION | T | | | |
| GOBP_DENDRITIC_SPINE_DEVELOPMENT | T | | | |
| GOBP_DENDRITIC_SPINE_MORPHOGENESIS | T | | | |
| GOBP_DETECTION_OF_STIMULUS_INVOLVED_IN_SENSORY_PERCEPTION | | | | VS |
| GOBP_DOPAMINE_SECRETION | T | | | |
| GOBP_DOPAMINE_TRANSPORT | T | | | |
| GOBP_DOPAMINERGIC_NEURON_DIFFERENTIATION | T | | | |
| GOBP_DORSAL_VENTRAL_PATTERN_FORMATION | T | | | |
| GOBP_FOREBRAIN_CELL_MIGRATION | T | | | |
| GOBP_GLIAL_CELL_DIFFERENTIATION | T | | | |
| GOBP_MAINTENANCE_OF_SYNAPSE_STRUCTURE | T | | | |
| GOBP_MEMORY | T | | | |
| GOBP_MORPHOGENESIS_OF_A_BRANCHING_STRUCTURE | T | | | |
| GOBP_MOTOR_NEURON_AXON_GUIDANCE | T | | | |
| GOBP_NEGATIVE_REGULATION_OF_AXON_EXTENSION | T | | | |
| GOBP_NEGATIVE_REGULATION_OF_NERVOUS_SYSTEM_DEVELOPMENT | T | | | |
| GOBP_NEGATIVE_REGULATION_OF_NEURAL_PRECURSOR_CELL_PROLIFERATION | T | | | |
| GOBP_NEURAL_PRECURSOR_CELL_PROLIFERATION | T | | | |
| GOBP_NEUROBLAST_PROLIFERATION | T | | | |
| GOBP_NEUROMUSCULAR_SYNAPTIC_TRANSMISSION | T | | | |
| GOBP_NEURON_PROJECTION_ORGANIZATION | T | | | |
| GOBP_OPTIC_NERVE_DEVELOPMENT | T | V | | |
| GOBP_POSITIVE_REGULATION_OF_NEURAL_PRECURSOR_CELL_PROLIFERATION | T | | | |
| GOBP_POSITIVE_REGULATION_OF_NEUROBLAST_PROLIFERATION | T | | | |
| GOBP_POSTSYNAPSE_ORGANIZATION | T | | | |
| GOBP_PRESYNAPSE_ORGANIZATION | T | | | |
| GOBP_PROTEIN_LOCALIZATION_TO_SYNAPSE | T | | | |
| GOBP_REGULATION_OF_DENDRITE_EXTENSION | T | | | |
| GOBP_REGULATION_OF_NEURAL_PRECURSOR_CELL_PROLIFERATION | T | | | |
| GOBP_REGULATION_OF_NEUROBLAST_PROLIFERATION | T | | | |
| GOBP_REGULATION_OF_NEUROTRANSMITTER_TRANSPORT | T | | | |
| GOBP_REGULATION_OF_POSTSYNAPTIC_MEMBRANE_POTENTIAL | T | | | |
| GOBP_SENSORY_ORGAN_MORPHOGENESIS | | V | | |
| GOBP_SENSORY_PERCEPTION | T | | | VS |
| GOBP_SENSORY_PERCEPTION_OF_CHEMICAL_STIMULUS | T | | | VS |
| GOBP_SENSORY_PERCEPTION_OF_PAIN | T | | | |
| GOBP_SENSORY_PERCEPTION_OF_SMELL | T | | | |
| GOBP_SENSORY_PERCEPTION_OF_TASTE | | | | VS |
| GOBP_SPINAL_CORD_DEVELOPMENT | T | | | VS |
| GOBP_SYNAPTIC_TRANSMISSION_GLUTAMATERGIC | | | T | |
| GOCC_ASTROCYTE_PROJECTION | T | | | |
| GOCC_AXONAL_GROWTH_CONE | T | | | |
| GOCC_EXCITATORY_SYNAPSE | T | | | |
| GOCC_GLIAL_CELL_PROJECTION | T | | | |
| GOCC_GLUTAMATERGIC_SYNAPSE | T | | | |
| GOCC_NEUROMUSCULAR_JUNCTION | | | | VS |
| GOCC_NEURON_PROJECTION_TERMINUS | T | | | |
| GOCC_NEURON_SPINE | T | | | |
| GOCC_NEURON_TO_NEURON_SYNAPSE | T | | | |
| GOCC_POSTSYNAPTIC_DENSITY_MEMBRANE | T | | | |
| GOCC_POSTSYNAPTIC_MEMBRANE | T | | | |
| GOCC_POSTSYNAPTIC_SPECIALIZATION | T | | | |
| GOCC_POSTSYNAPTIC_SPECIALIZATION_MEMBRANE | T | | | |
| GOCC_PRESYNAPTIC_MEMBRANE | T | | | |
| GOCC_SYNAPTIC_MEMBRANE | T | | | |
| GOMF_ACETYLCHOLINE_RECEPTOR_ACTIVITY | | V | | |
| GOMF_G_PROTEIN_COUPLED_SEROTONIN_RECEPTOR_ACTIVITY | | V | | |
| GOMF_GLUTAMATE_RECEPTOR_ACTIVITY | T | | | |
| GOMF_NEUROTRANSMITTER_RECEPTOR_ACTIVITY | T | V | | |
| GOMF_POSTSYNAPTIC_NEUROTRANSMITTER_RECEPTOR_ACTIVITY | T | | | |

**Table S3.** List of enriched neuronal genesets from the ‘C5’ group (GENE ONTOLOGY) of curated genesets from Molecular Signatures Database (MSigDB) obtained from GSEA. The denoted condition under each pairwise comparison for each geneset represents the enriched condition. They are called as significant by both RNA-seq pipelines.

| Autophagy geneset | Stringtie | feature Counts |
|---|---|---|
| GOBP_AUTOPHAGY_OF_MITOCHONDRION | TS, VS | VS |
| GOBP_MACROAUTOPHAGY | VS | TS, VS |
| GOBP_NEGATIVE_REGULATION_OF_AUTOPHAGY | | TS, VS |
| GOBP_REGULATION_OF_AUTOPHAGY_OF_MITOCHONDRION | TS, VS | TS, VS |
| KEGG_MEDICUS_REFERENCE_AUTOPHAGY_VESICLE_NUCLEATION_ELONGATION_MATURATION_MTORC1_PI3KC3_C1 | TS | TS |
| WP_HOST_PATHOGEN_INTERACTION_OF_HUMAN_CORONAVIRUSES_AUTOPHAGY | TS, VS | |
| WP_PERTURBATIONS_TO_HOST_CELL_AUTOPHAGY_INDUCED_BY_SARS_COV_2_PROTEINS | TS | TS |

**Table S4.** List of enriched autophagy datasets in the denoted starvation conditions compared to their respective fed conditions ('V vs VS' and 'T vs TS') from GSEA.

| | |
|---|---|
| **ATGs** | ATG4A, ATG4B, ATG4C, ATG5, ATG12, ATG16L2, ATG3, ATG7, ATG101, ATG2A, ATG2B, ATG14, ATG9A, ATG9B, ATG13, ATG10, ATG8s (MAP1LC3A, MAP1LC3B, GABARAP, GABARAPL1, GABARAPL2), ULK1, ULK2, ULK3, BECN1, BCL2, VMP1, AMBRA1, UVRAG, ZFYVE1, TFs (TFEB, FOXO1, FOXO3), PINK1, MTOR, PRKAA1, PRKAA2, PIK3C3, PIKFYVE, SNAP29, SNX18, STX17, VAMP7, VAMP8, LAMP1, LAMP2 |
| **Endolysosomal channels** | TRPs (MCOLN1, MCOLN2, MCOLN3, TRPV4, TRPV6, TRPM2, TRPC1), TPCN1, TPCN2, KCNMA1, P2RX4, PACC1, TMEM175, HVCN1, VGSCs (SCN1A), SCNN1A, CLCN2, CLCN3, CLCN4, CLCN6, CLCN7, VRACs (LRRC8A, LRRC8B, LRRC8C, LRRC8D, LRRC8E), VGCCs (CACNA1A, CACNA2D1, CACNA1C, CACNB1), VGKCs (KCNJ3, KCNJ4, KCNJ8, KCNK3, KCNK6, KCNH2, KCNQ1), CFTR |
| **V-ATPase subunits** | ATP6V0A1, ATP6V0A2, ATP6V0B, ATP6V0C, ATP6V0D1, ATP6V0E1 |

**Table S5.** List of the 90 genes, classified into three categories, chosen for meta-analysis. *TF – Transcription Factor, TRP – Transient Receptor Potential Channel, VGCC – Voltage Gated Calcium Channel, VRAC – Volume Regulated Anion Channel, VGSC – Voltage Gated Sodium Channel, VGKC – Voltage Gated Potassium Channel*

| Autophagy | | | | | Non-Autophagy | | | | |
|---|---|---|---|---|---|---|---|---|---|
| **PC1** | **PC2** | **PC3** | **PC4** | **PC5** | **PC1** | **PC2** | **PC3** | **PC4** | **PC5** |
| KCNJ3 | SCNN1A | VAMP8 | KCNJ8 | CACNA1C | LRRC8E | TRPC1 | TRPC1 | FOXO1 | CACNA2D1 |
| KCNJ4 | ULK1 | BCL2 | LRRC8C | KCNQ1 | VMP1 | VAMP8 | MCOLN3 | KCNMA1 | MCOLN3 |
| GABARAPL2 | LRRC8B | CFTR | MCOLN2 | MCOLN3 | PRKAA2 | TRPM2 | MAP1LC3B | ATG9B | KCNJ8 |
| CACNA2D1 | KCNQ1 | CACNA1C | ATG9B | SCNN1A | MAP1LC3B | TFEB | CLCN3 | SCN1A | CACNA1C |
| MAP1LC3A | ATG16L2 | ATP6V0B | CFTR | HVCN1 | ATP6V0D1 | BCL2 | GABARAPL2 | TRPC1 | CLCN4 |

**Table S6.** Top 5 genes in the first 5 Principal Components of both dataset groups

| **Sr. No.** | **Target** | **Primer Sequences (5'-3')** | **Amplicon Size** (bp) |
|---|---|---|---|
| 1 | TRPML3 | TGAGAAGTTCTGGGCTCGAG | 190 |
| | | TGTGTCATCCATTCGGTCCA | |
| 2 | BECN1 | GACACTCAGCTCAACGTCAC | 184 |
| | | CTGCCACTATCTTGCGGTTC | |
| 3 | FOXO1 | GCGCTTAGACTGTGACATGG | 153 |
| | | ACTAACCCTCAGCCTGACAC | |
| 4 | FOXO3 | CCAGCCTGTCACCTTCAGTA | 237 |
| | | TAAAGGAGCTGGTTGGGGAG | |
| 5 | LC3B | CACTCTTGCAGGGGTAGACA | 241 |
| | | TGTGTTTCTCCCGCTGTACT | |
| 6 | ATG12 | CTGGAGGGGAAGGACTTACG | 202 |
| | | AGTCCTTGGATGGTTCGTGT | |
| 7 | GABARAPL1 | TAGATGGGTCAGGAGGTGGA | 250 |
| | | CTGAGTGAGATTGCAACCGG | |
| 8 | GABARAPL2 | CCAGCCAATTCATGAGTCGG | 195 |
| | | TGCAGCAAGACCTACATCCA | |
| 9 | TFEB | CCATCCCCATTCCATCACCT | 193 |
| | | ACAGAAGTGGATCAGAGGCC | |
| 10 | LAMP2 | TCTAGAGGTTAGTGGGCCCT | 183 |
| | | CCAATCAAGCAGACACAGGG | |
| 11 | ULK1 | GACCGCATTCACAGCATCAC | 87 |
| | | GAACATCTCGTCCAGGGCAG | |
| 12 | ATG9A | CCTCATCCTCATCTTCTGCCTGC | 92 |
| | | TATCTCCCACACCAACGACCTCC | |
| 13 | LRRC8C | CTGCCTCCACCTAAACCATCTC | 119 |
| | | AGGGCTCGCTCATAACACATC | |
| 14 | KCNJ3 | GGCAGCGGTTCGTGGACAAG | 150 |
| | | CGGTGTAGGTGAGAATGAAGATGAAGAG | |
| 15 | B-Actin | CATCCGCAAAGACCTGTACG | 218 |
| | | CCTGCTTGCTGATCCACATC | |
| 16 | GAPDH | CACATCGCTCAGACACCATG | 198 |
| | | TGACGGTGCCATGGAATTTG | |

**Table S7.** Details of the gene targets probed by qPCR, their primer sequences and amplicon sizes
